# The Trouble with Triples: Examining the Impact of Measurement Error in Mediation Analysis

**DOI:** 10.1101/2022.07.07.499004

**Authors:** Madeleine S. Gastonguay, Gregory R. Keele, Gary A. Churchill

## Abstract

Mediation analysis is used in genetic mapping studies to identify candidate gene mediators of quantitative trait loci (QTL). We consider genetic mediation analysis of triplets - sets of three variables consisting of a target trait, the genotype at a QTL for the target trait, and a candidate mediator that is the abundance of a transcript or protein whose coding gene co-locates with the QTL. We show that, in the presence of measurement error, mediation analysis can infer partial mediation even in the absence of a causal relationship between the candidate mediator and the target. We describe a measurement error model and a corresponding latent variable model with estimable parameters that are combinations of the causal effects and measurement errors across all three variables. The relative magnitudes of the latent variable correlations determine whether or not mediation analysis will tend to infer the correct causal relationship in large samples. We examine case studies that illustrate the common failure modes of genetic mediation analysis and demonstrate how to evaluate the effects of measurement error. While genetic mediation analysis is a powerful tool for identifying candidate genes, we recommend caution when interpreting mediation analysis findings.

## Introduction

Mediation analysis is a class of statistical methods used to determine whether the effect of an exogenous variable (*X*) on a target (*Y*) may be wholly, partially, or not transmitted through a mediator (*M*) (Figure 1). We want to determine which of the causal effects, represented by edges in the graph, are present or absent. For example, if the effect of *X* on *Y* is wholly mediated through *M*, the edge labeled *c* is absent, indicating that there is no direct causal effect of *X* on *Y*. Mediation inference relies on propagation of variation between causally linked variables that produces characteristic patterns of correlation in data. However, in addition to causal variation, data often include a measurement error component that does not propagate. In many applications of mediation analysis, measurement error is not accounted for and the resulting model mis-specification can potentially bias mediation inference (Richmond *et al*. 2016).

**Figure 1.**
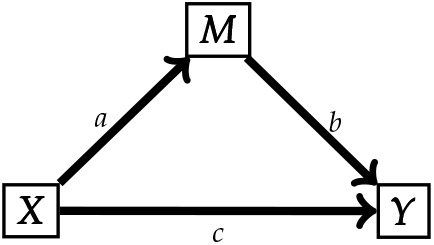
A simple mediation model. *X* effects *Y* directly (*c*) and indirectly through *M* (*ab*).

Commonly used methods for mediation inference such as causal steps (Baron and Kenny 1986) and the Sobel test (Sobel 1982) seek to establish the presence of an indirect effect of *X* on *Y* through *M*, *i.e*., edges labeled *a* and *b* are both present. Here, we apply a more general Bayesian model selection approach to infer the most likely causal structure relating *X*, *M*, and *Y* (Crouse *et al*. 2022). Importantly, our findings are not dependent on the inference method as they follow from properties of the underlying model. Bayesian model selection is likelihood-based; it performs as well or better than other inference methods. Unlike the Sobel test, Bayesian model selection distinguishes complete from partial mediation. However, as with other commonly used mediation analysis methods, it does not account for measurement error.

Previous studies of the impact of measurement error on mediation analysis have demonstrated bias in estimation of the direct effect (the effect of *X* on *Y* independent from *M*) (le Cessie *et al*. 2012). In addition, measurement error in *M* results in underestimation of the indirect effect (the effect of *X* on *Y* through *M*), and loss of conditional independence between *X* and *Y*, even when the effect of *X* on *Y* is fully mediated through *M* (Ledgerwood and Shrout 2011; Otter *et al*. 2018; Pierce *et al*. 2014; Rockman 2008; VanderWeele *et al*. 2012). These impacts are similar to those that arise with un-modeled confounders (Fritz *et al*. 2016; Liu and Wang 2021) and other generalizations of the three-variable mediation model (Cole and Preacher 2014). It is known that unaccounted measurement error can lead to inference of partial mediation even when there is no direct effect on *Y* through *M* (Otter *et al*. 2018; Pierce *et al*. 2014; Gonzalez and MacKinnon 2021), a point we emphasize here.

Our aim in this work is to assess the impact of measurement error on mediation analysis in genetic mapping studies where *X* is the genotype at a quantitative trait locus (QTL) associated with a target phenotype *Y*, and the candidate mediator *M* is the expression level of a transcript or protein product of a gene that co-localizes with the QTL, *i.e*., a local gene expression or protein QTL (eQTL or pQTL). Genetic mediation analysis was introduced by Schadt *et al*. (2005) and has been widely adopted in various forms in model organism genetic mapping studies (e.g., Keller *et al*. (2018)). More recently, it has been applied in transcript-wide association studies (TWAS) (Li and Ritchie 2021) to identify gene expression mediators of clinical traits in humans. We caution that co-localization of the target trait QTL and the candidate mediator is not sufficient to establish a causal relationship because an association between *M* and *Y* could result from linked genetic variants with unrelated causal effects; it could be induced by an unobserved confounder of *M* and *Y*, a problem that Mendelian randomization (MR) is designed to address (Katan 1986; Didelez and Sheehan 2007); or, as we demonstrate here, it could be an artifact due to measurement error.

## Results

### Bayesian Model Selection

Our objective is to determine the structure of the causal relationships among *X*, *M*, and *Y*. Specifically, we want to determine if one or more of the edges in Figure 1 is absent. We adopt terminology from genetic mediation analysis (Schadt *et al*. 2005; Neto *et al*. 2013) and refer to the causal structures of interest as Causal (*c* = 0), Independent (*b* = 0), Reactive (*a* = 0), and Complex (all effects are non-zero). In addition, there are causal structures for which the three variables are not fully connected that we refer to as non-mediation models. Under the Causal model, also known as complete mediation, the effects of *X* on *Y* are completely mediated through *M*. Under the Independent model, *X* has direct but independent effects on each of *M* and *Y*. Under the Reactive model, the roles of the mediator and target variables are reversed such that *M* is responding to variation in *Y*. Finally, under the Complex model, also known as partial mediation, *X* affects *Y* directly and indirectly through *M*. We note that there are challenges in distinguishing the directionality of the relationship between *M* and *Y* (Wiedermann and von Eye 2015). In some cases, the context will determine if causation from *Y* to *M* is possible.

Bayesian model selection as implemented in the bmediatR software (Crouse *et al*. 2022) provides a likelihood-based decision rule that selects the model with the highest posterior probability among a predefined set of models. It does not rely on hypothesis testing and thus avoids the difficulties inherent in establishing a null hypothesis, *i.e*., an effect size of zero. We applied bmediatR with a uniform prior across the set of models and weakly informative priors on the effect sizes. We either restrict model selection to choose among the Causal, Independent, and Reactive models or we consider an expanded set of models that includes Complex and other non-mediation alternatives. As noted above, the model likelihood implemented in bmediatR does not account for measurement error.

### Measurement Error Models

To incorporate measurement error, we introduce the error-free values of the *causal variables*, denoted as *X*\*, *Y*\*, and *M*\*. The causal variables are not directly observed. Instead we observe their surrogates, the *measured variables X*, *Y*, and *M*. We define the measurement error model in terms of correlation parameters (Figure 2A; Appendix). The *causal correlations* (*ρ_X*M*_*, *ρ_X*Y*_*, and *ρ_Y*M*_*) determine the relationship among the causal variables, which we refer to as the *causal structure* of the measurement error model. The *error correlations* (*ρ_X*X_*, *ρ_Y*Y_*, and *ρ_M*M_*) determine the level of measurement error between each causal variable and its measured counterpart. The use of error correlations, which are equivalent to but inversely related to the more conventional error variances, simplifies our specification of the measurement error model. We assume that measurement error is independent for each pair of variables and that there are no hidden confounders. While these are strong assumptions, incorporating these features into our model would only further obstruct our ability to infer the correct causal structure.

**Figure 2.**
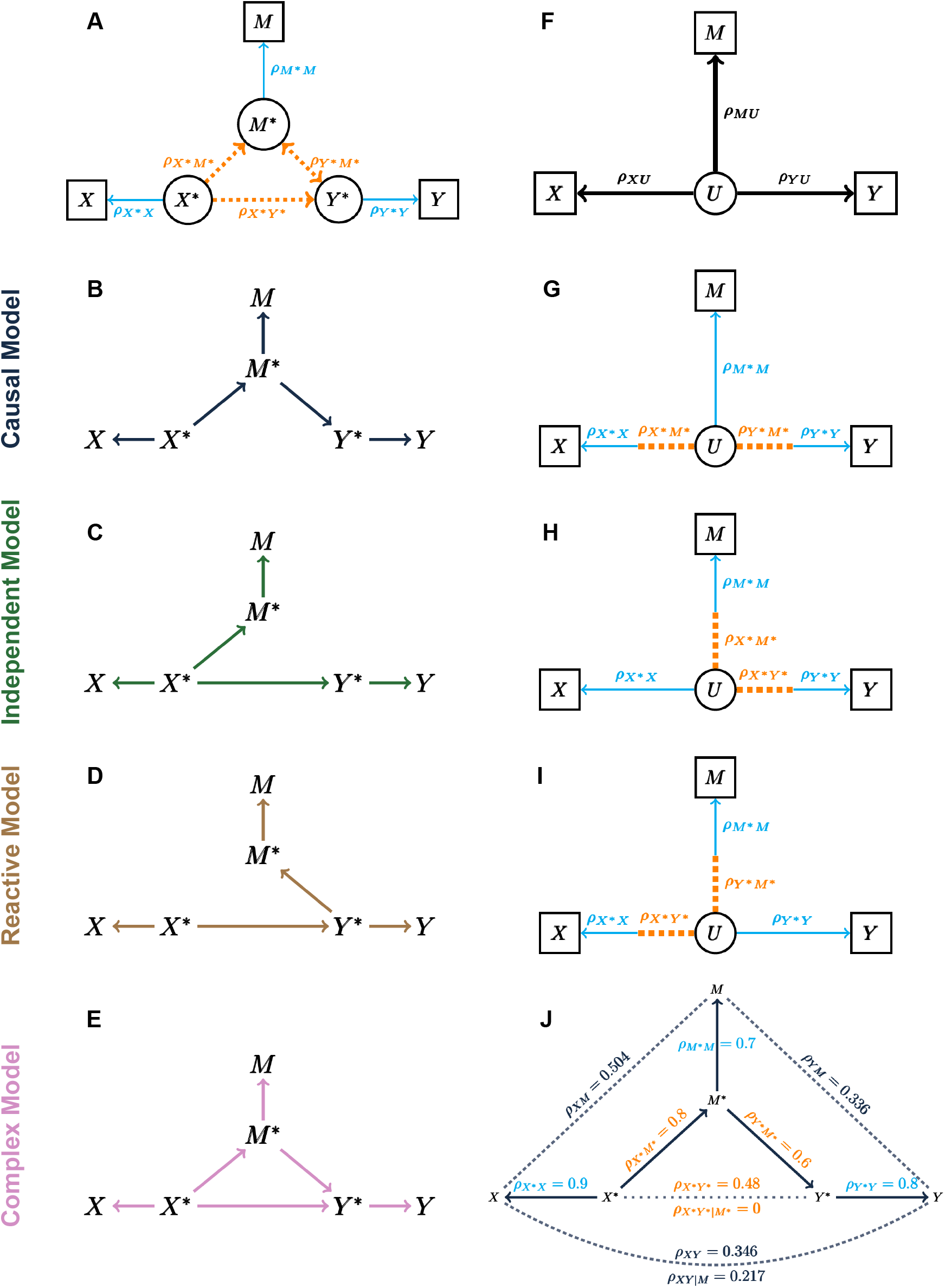
Directed acyclic graphs (DAGs) of measurement error models and their corresponding latent variable models. (A) The measurement error model describes the relationships among causal variables in terms of their causal correlations (dotted orange lines) and between each causal variable and its corresponding measured variable in terms of error correlations (solid blue lines). Variables in circles are unobserved and those in boxes are measured. (B-E) DAGs for the Causal, Independent, Reactive, and Complex measurement error models. (F) The structure of the latent variable model where *U* is a single unobserved (latent) variable. (G-I) The latent correlations are determined by the causal correlations (dotted orange edges) and error correlations (solid blue edges) in different combinations for the Causal, Independent, and Reactive models, respectively. (J) An example of the Causal measurement error model for the Causal model. Causal (partial) correlations are labeled in orange, error correlations are labeled in bright blue, and data (partial) correlations are labeled in dark blue. Dotted lines denote correlations between variables that do not share a direct edge in the measurement error model.

We define the *data correlations* (*ρ_XY_*, *ρ_XM_*, and *ρ_YM_*) to be the expected correlations among the measured variables. They can be expressed in terms of the causal and error correlations:

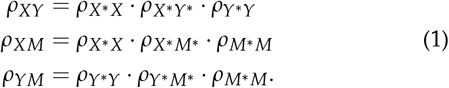

The data correlations are always weaker than their corresponding causal correlations, more so when there is more measurement error. The three data correlations can be estimated from observed data, but we cannot estimate the causal and error correlations without additional information or constraints (see Appendix).

The Causal, Independent, and Reactive structures (Figure 2B-E) impose constraints on the causal correlations. It is convenient here to introduce the term *middle variable*. For the Causal model, the middle variable is M, for the Independent model, it is X, and for the Reactive model, it is Y. In each case, the two non-middle variables are not directly connected by an edge. The causal correlation between the two non-middle variables is constrained to equal the product of the other two causal correlations. Equivalently, their partial correlation after conditioning on the middle variable is equal to zero. For the Complex model, the causal correlations are not constrained, aside from the requirement that the correlation matrix is positive semi-definite.

Consider the Causal measurement error model with parameters as specified in Figure 2J. The causal correlations satisfy the constraint *ρ_X*Y*_* = *ρ_X*M*_* · *ρ_Y*M*_* or equivalently, *ρ_X*Y*|M*_* = 0. However, the data correlations do not satisfy these constraints; even a small amount of measurement error can result in data correlations that differ substantially from the causal correlations. The key contributor to this discrepancy is the error correlation of the middle variable, in this example *ρ_M*M_*. The data correlations will satisfy the same constraints as the causal correlations if and only if there is no measurement error in the middle variable (see Appendix). In the presence of measurement error on the middle variable, the data correlations will be unconstrained as they are for the Complex model without measurement error.

In summary, data correlations among measured variables *X*, *M*, and *Y* need not satisfy the constraints implied by the causal structure. Indeed, all forms of the measurement error model (Causal, Reactive, Independent, and Complex) are likelihood equivalent to the Complex model without measurement error (see Appendix). This presents a dilemma for determining the causal structure from real data. Of course, estimated data correlations will never exactly satisfy these constraints and statistical inference is needed to determine if the observed data are consistent with an assumed model. Before we turn to the question of whether and when it is possible to recover the correct causal structure from observed data in the presence of measurement error, we introduce a simplified form of the measurement error model.

### The Latent Variable Model

The Causal, Independent, and Reactive measurement error models have five free parameters (six parameters with one constraint) and three observable outcomes. We can transform each of these models to an equivalent latent variable model with three free parameters. In the latent variable model, the causal variables *X*\*, *M*\*, and *Y*\* are replaced by a single *latent variable, U* (Figure 2F). The *latent correlations* (*ρ_XU_*, *ρ_YU_*, and *ρ_MU_*) determine the relationship between *U* and each of the measured variables, and the data correlations can be expressed in terms of latent correlations:

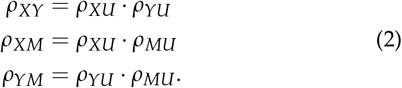

We can also express the latent correlations in terms of the causal and error correlations (Table 1, Figure 2G-I). The expressions depend on which causal structure is assumed. For the non-middle variables, the latent correlations are the product of a causal correlation and an error correlation, and for the middle variable, the latent correlation is equal to its error correlation. The latent variable model parameters are estimable from data (see Appendix). Knowing which (products of) parameters can be estimated for a given causal structure, will be helpful when diagnosing the impacts of measurement error.

**Table 1.** Latent Correlations for each model.

| Causal Structure | Latent Correlation |  |  |
| --- | --- | --- | --- |
| | $\rho_{MU}$ | $\rho_{XU}$ | $\rho_{YU}$ |
| Causal | $\rho_{M^*M}$ | $\rho_{X^*X} \cdot \rho_{X^*M^*}$ | $\rho_{Y^*Y} \cdot \rho_{Y^*M^*}$ |
| Independent | $\rho_{M^*M} \cdot \rho_{X^*M^*}$ | $\rho_{X^*X}$ | $\rho_{Y^*Y} \cdot \rho_{X^*Y^*}$ |
| Reactive | $\rho_{M^*M} \cdot \rho_{Y^*M^*}$ | $\rho_{X^*X} \cdot \rho_{X^*Y^*}$ | $\rho_{Y^*Y}$ |

### Simulations

The parameters of the measurement error model, denoted as *ρ*, can be thought of as correlations estimated from infinitely large data. We now consider what happens when model structure is inferred from correlations estimated from data from finite samples, denoted as *r*.

We simulated data from the measurement error model and then analyzed the data assuming no measurement error. Specifically, we simulated *X*\* from a standard normal distribution and then simulated *M*\*, *Y*\*, *X*, *M*, and *Y* according to linear models with the desired causal or error correlations (Crouse *et al*. 2022) (see Methods). The sample size for simulated data ranged from *N* = 200 to *N* = 5000. We simulated 100,000 data sets for each of the Causal, Independent, and Reactive measurement error models. Model parameters (causal and error correlations) were sampled independently from beta distributions to obtain data correlations similar to those that arise in practice. We used bmediatR (Crouse *et al*. 2022) to select the model with the greatest posterior probability for each simulated data set. We first restricted the model selection to choose among only the Causal, Independent, or Reactive models (Schadt *et al*. 2005; Neto *et al*. 2013) (*three-choice model options*). We then repeated the model selection including the Complex model and non-mediation models (*expanded model options*).

Our rationale for examining three-choice model selection is partly motivated by previous approaches to genetic mediation analysis (Schadt *et al*. (2005)). In addition, recognizing that the measurement error model is equivalent to partial mediation (Complex model), we expect that Bayesian model selection in large samples will infer the Complex model regardless of the underlying causal structure. In the simulations and data examples presented below we apply both options.

#### Three-choice model selection

The estimated data correlations obtained from simulations of the Causal, Independent, and Reactive measurement error models have overlapping ranges, *i.e*., a large proportion of the estimated data correlations could have been obtained from any of the three causal structures (Figure 3). This immediately suggests that it will be difficult to distinguish among these models. Overall, we correctly classified the causal structure for ~ 62% of simulated data sets with *N* = 200 (Table 2). The rate of correct classification across all parameter configurations increased with increasing sample size, but never exceeded 65% for sample sizes up to *N* = 5000 (Figure S1).

**Figure 3.**
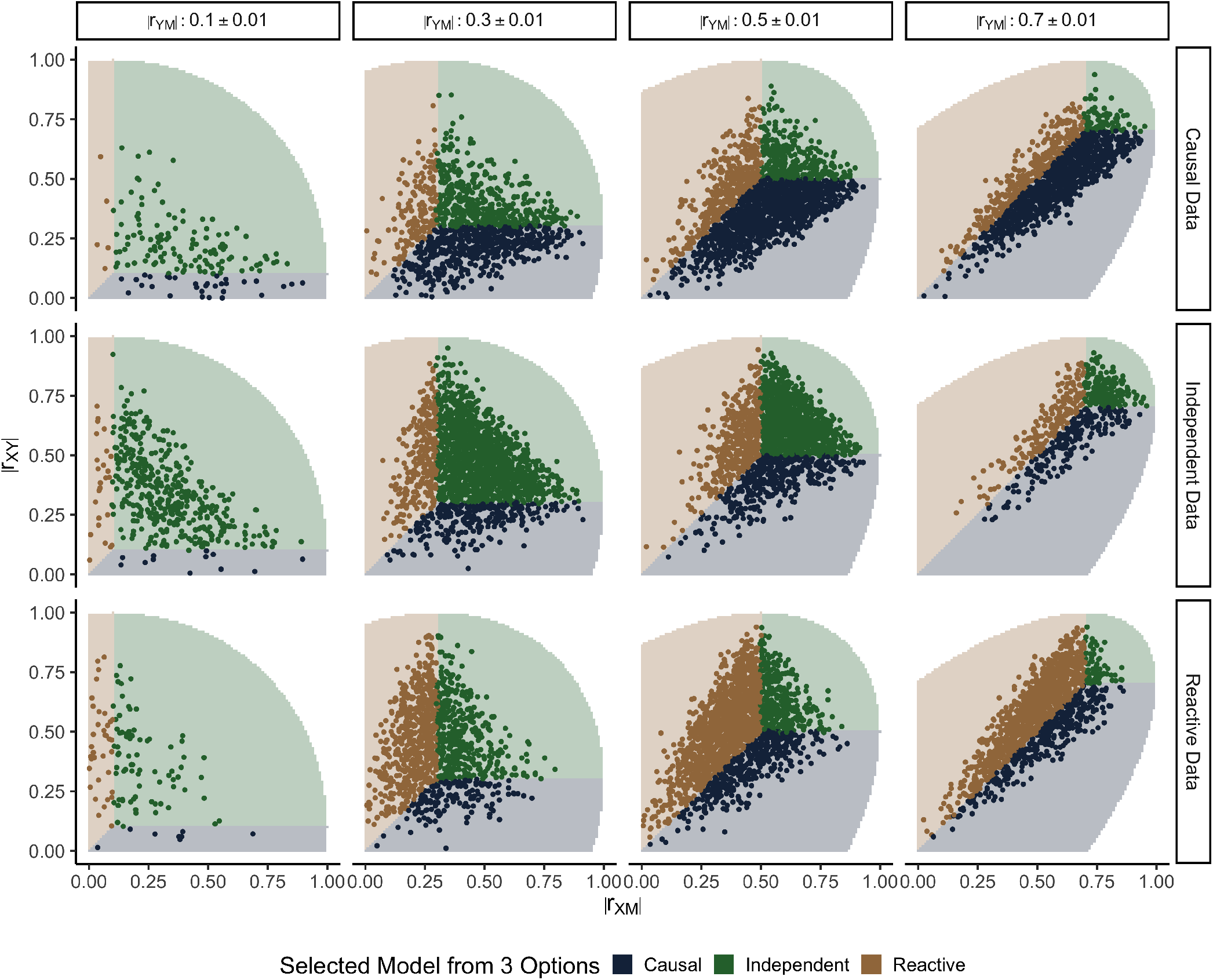
Three-choice model selection outcomes are determined by the estimated data correlations. Each row of panels corresponds to simulations of a different causal structure (*N* = 200). Columns correspond to binned values of *r_YM_*. The x- and y-axes show *r_XM_* and *r_XY_*, respectively. Points representing the estimated data correlations are colored to indicate the model with the greatest posterior probability from three-choice model selection. Shaded regions indicate the range of data correlations in which each model will be inferred, and the unshaded region delineates where the correlation matrices are not positive semi-definite. Table 2 summarizes the model selection outcomes over all simulated parameter settings

**Table 2.**
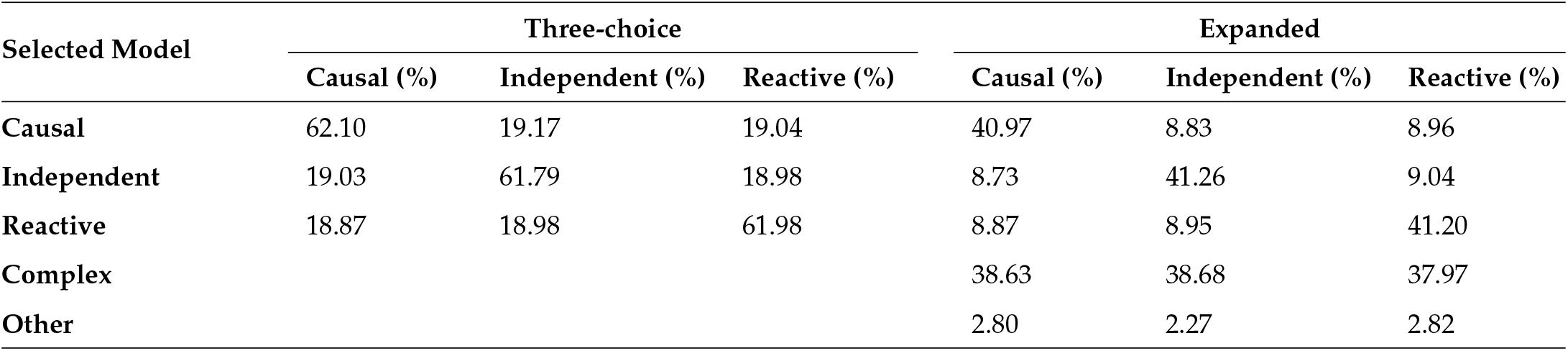
Classification of simulated data across all sampled parameters for three-choice and expanded model selection (*N* = 200).

Bayesian three-choice model selection always selected the model that has no direct effect between the two causal variables whose estimated data correlation is the weakest (see shading in Figure 3). For example, if the correlation between *X* and *Y* is less (in magnitude) than the other two data correlations, the Causal model will be selected. This simple inference rule for three-choice model selection is useful but it only holds for univariate *X*, *M*, and *Y*.

#### Expanded model selection

When we expanded the model selection options to include the Complex and non-mediation models, we saw a decrease in the overall rate of correct classification (41% when *N* = 200) but an even greater proportional decrease in the rate of incorrect classification as one of the constrained models (9% when *N* = 200, Table 2). When *N* = 200, the Complex model was selected almost as frequently as the correct causal structure and for larger sample sizes, the rate of selecting the Complex model increased, *e.g*., up to 88% when *N* = 5000 (Figures S1 and S2).

When using the expanded model selection options, the simple inference rule (shading) in Figure 4 no longer applied, but there were some regularities. The Complex model was selected when all three data correlations were similar in magnitude. A non-mediation model was selected when at least two of the data correlations were sufficiently weak. However, if one of the three constrained models was selected, it conforms with the inference rule from three-choice model selection.

**Figure 4.**
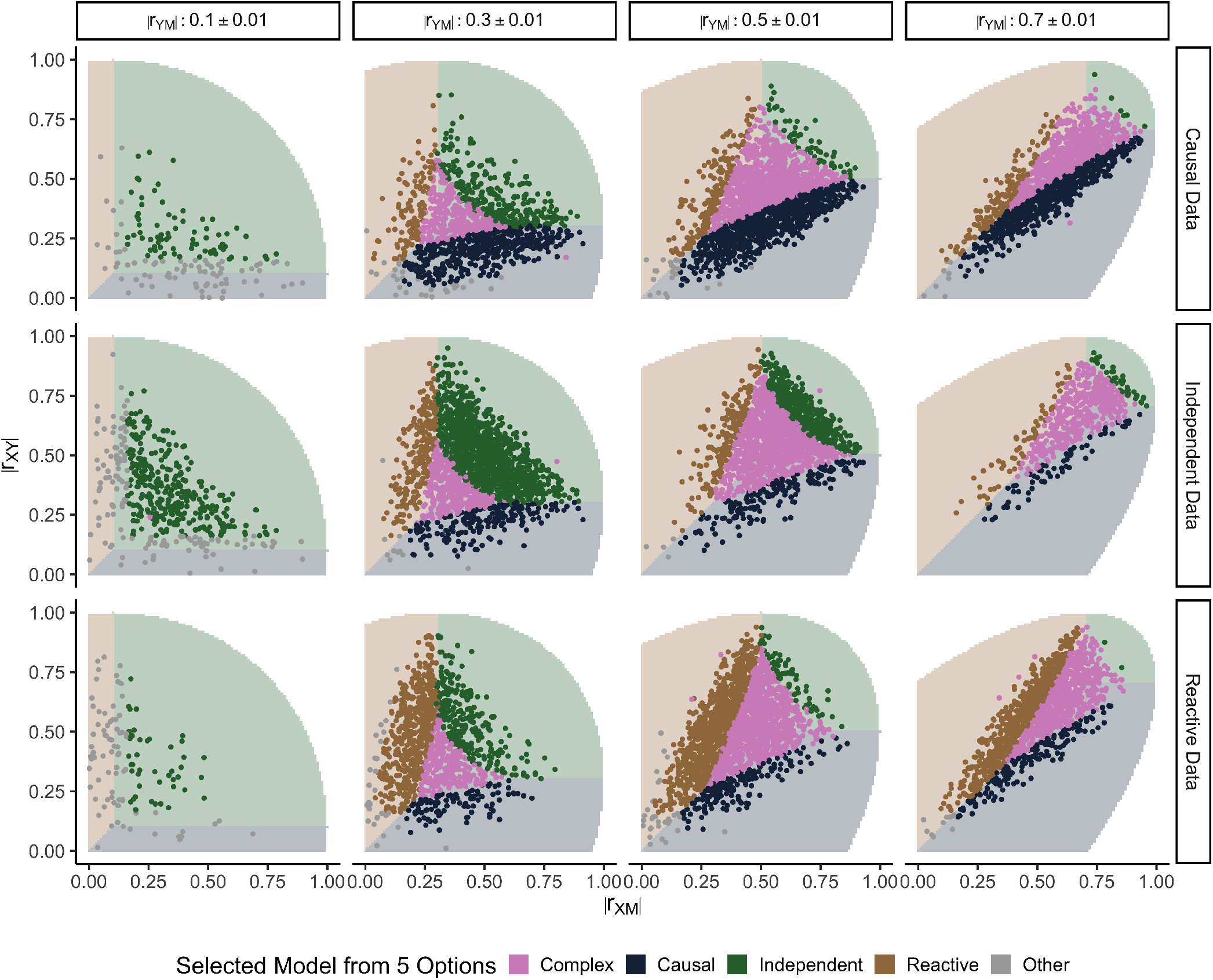
Expanded model selection outcomes as a function of estimated data correlations. Each row of panels corresponds to simulations of a different causal structure (*N* = 200). Columns correspond to binned values of *r_YM_*. The x- and y-axes show *r_XM_* and *r_XY_*, respectively. Points representing the estimated data correlations are colored to indicate the model with the greatest posterior probability from expanded model selection. Shaded regions indicate the three-choice model selection inference rule, and the unshaded region delineates where the correlation matrices are not positive semi-definite. Table 2 summarizes model selection outcomes over all simulated parameter settings *Y* and *M*.

#### Latent correlations determine the consistency of mediation analysis

The asympototic behavior of three-choice model selection is determined by the latent correlations (Table 1). For example, if *ρ_MU_* is the strongest latent correlation, then *ρ_XY_* will be the weakest data correlation (Equation 2). As sample size increases, the estimated latent correlation *r_MU_* will converge towards its true value and, applying the simple inference rule, three-choice model selection will tend toward selecting the Causal model. Similarly, when *ρ_XU_* or *ρ_YU_* are the strongest latent correlations, the Independent and Reactive models will be selected more frequently as sample size increases, respectively. Thus, it is the relative sizes of the latent correlations that determine whether three-choice Bayesian model selection will be consistent (tending toward the correct causal structure) or inconsistent (tending toward an incorrect causal structure). This is confirmed in simulations of the Causal model with increasing sample size (Figure 5A).

**Figure 5.**
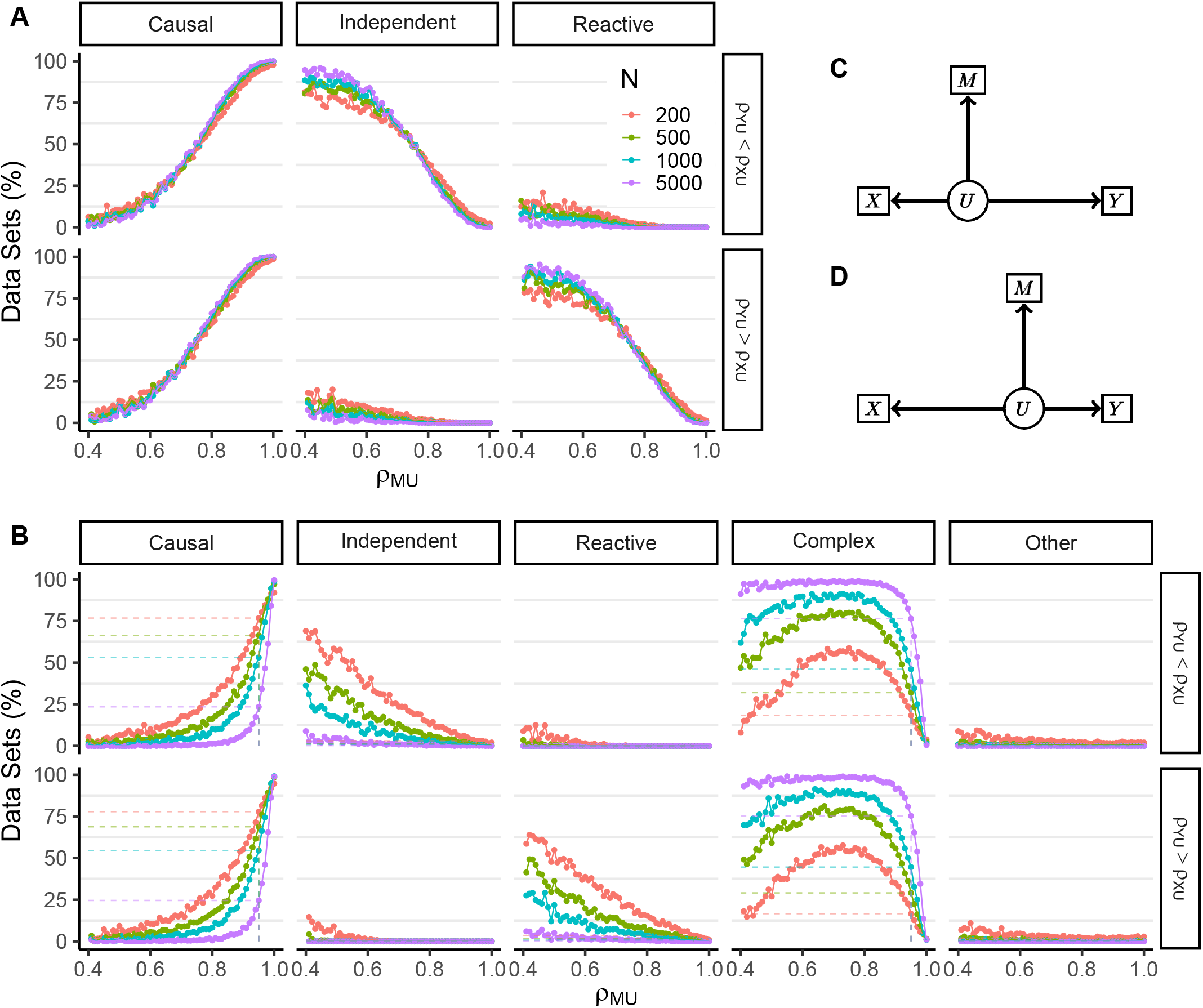
Classification rates as a function of error in *M* for data simulated from the Causal measurement error model. Classification rates are shown for three-choice (A) or expanded (B) model options. Causal classifications are correct and non-Causal classifications are incorrect. Line color denotes sample size used in simulations. Each column shows the percent of data sets classified as the model listed at the top as a function of the latent correlation *ρ_MU_*. Dashed lines in (B) mark results when *ρ_MU_* = 0.95. Both (A) and (B) are split into two rows showing results for data simulated with a stronger latent correlation for *X* (top row), and data sets with a stronger latent correlation for *Y* (bottom row). The corresponding latent variable DAGs are displayed in (C) and (D) where shorter edges denote stronger correlations.

The latent correlations also determine the large-sample behavior of Bayesian model selection with expanded model options (Figure 5B). We found that for simulations of the Causal measurement error model, a strong *ρ_MU_* was required to drive selection of the Causal model. The Complex model was frequently selected at weaker values of *ρ_MU_*, regardless of the strength of *ρ_XU_* and *ρ_YU_*. Even with very small error in the mediator (*ρ_M*M_* = 0.95), the Causal model was selected for only 25% of data sets simulated at *N* = 5000. The remaining data sets were classified as Complex. Similar results were obtained for the Independent and Reactive models. Mediation analysis tends to support partial mediation when there is any measurement error in the middle variable. This should not be not surprising, as all forms of the measurement error model are likelihood equivalent to the Complex model without measurement error. Thus, selection of the Complex model is expected, but it is not useful for determining the causal structure.

#### Using latent correlations to diagnose the outcome of mediation analysis

The latent model determines which incorrect inferences are most likely. For data generated from a Causal model, three-choice model selection will be consistent if *ρ_MU_* is the strongest latent correlation; it will be inconsistent in the direction of the Reactive model if *ρ_YU_* is the strongest; and it will be inconsistent in the direction of the Independent model if *ρ_XU_* strongest.

We cannot directly estimate the causal and error correlations, but we can place constraints on their values (Figures 6, S3, and S4). To illustrate, suppose that three-choice model selection indicates the Causal model. We can evaluate how each of the possible causal structures could have given rise to this outcome. If the true causal structure is Causal, the measurement error model parameters will be consistent with one of the DAGs in Figure 6A. There may be equal error in all three variables; there may be less error in the mediator than the other variables; or there may be more error in the mediator than the other variables, but the causal correlations are weak. If the true causal structure is Independent, there must be more error in the exogenous variable than in the candidate mediator and *X*\* should be tightly correlated with *M*\* (Figure S3B). Lastly, if the true causal structure is Reactive, the measurement error for the target must be greater than for the mediator and *Y*\* is strongly correlated with *M*\* (Figure S4C). Ironically, weaker causal correlations and more measurement error in the non-middle variables can improve our ability to select the correct casual structure. A complete enumeration of scenarios that lead to consistent or inconsistent inferences using three-choice model selection is provided in Table S1.

**Figure 6.**
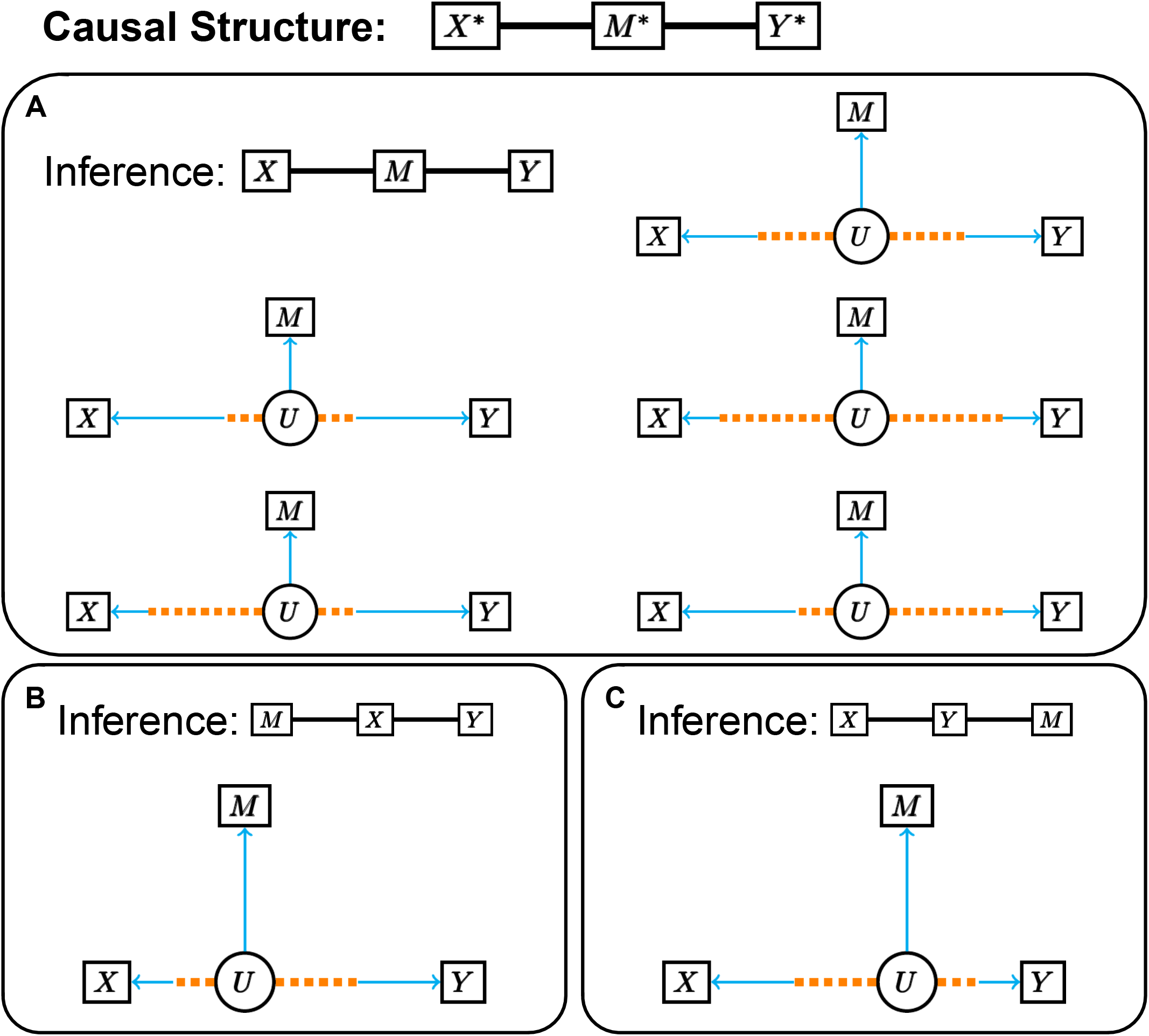
Latent variable representation of the Causal measurement error model. Configurations of the latent variable model representing the Causal model that result in (A) consistent and (B-C) inconsistent inferences. Blue edges correspond to the proportion of the latent correlation determined by the error correlation and dotted orange edges correspond to the proportion determined by the causal correlation. Shorter edges represent stronger correlation. (A) The correct model is inferred if the latent correlation for *M* is the strongest (the latent variable arm for *M* is the shortest). This can be achieved if there is an equal amount of error in all three variables (top right) or if there is less error in *M* than *X* and *Y* (middle left). If *M* is noisier than *X* and *Y*, the correct model may still be inferred if the causal correlations are weak (middle right). The bottom row shows scenarios where *X* and *Y* satisfy different configurations. (B) The Independent model is inferred if the latent correlation for *X* is the strongest. When the causal structure is the Causal model, this will only occur if the error correlation for *M* is weaker than both the error correlation and causal correlation contributing to the latent correlation for *X*. The composition of the latent correlation arm for *Y* does not influence the inference. (C) Shows the analogous scenario to (B) for inferring the Reactive model by swapping *X* and *Y*.

### Evaluating mediation analysis with real data

We obtained transcript and protein profiling data from liver tissue for 192 Diversity Outbred (DO) mice, including mice of both sexes (Chick *et al*. 2016), and for 116 Collaborative Cross (CC) mice, with one female and one male from each of 58 CC strains (Keele *et al*. 2021). The DO mice are an outbred stock, of which each mouse is a genetically unique individual and predominantly heterozygous at loci across the genome (Churchill *et al*. 2012). The CC mice represent a panel of recombinant inbred strains and are homozygous across most of their genomes(Collaborative Cross Consortium 2012; Srivastava *et al*. 2017). The DO and CC mice are descended from the same eight founder strains and they share the same genetic variants. To carry out genetic mediation analysis, we represented the genotype of each animal as an eight-state vector of haplotype dosages (Gatti *et al*. 2014), *i.e., X* is a multi-state exogenous variable.

For each study, we identified genes with local protein abundance QTL (pQTL) and a corresponding local gene expression QTL (eQTL). We refer to these as *concordant triplets* (genotype-transcript-protein), and we assume that in most cases the transcript will mediate the effect of genetic variation on protein abundance (Chick *et al*. 2016). We found 2023 concordant triplets in the DO data, 967 in the CC data, and 582 genes with concordant triplets in both studies. For each concordant triplet, we identified a transcript from a nearby gene with the strongest co-mapping local eQTL within 1Mb of the pQTL. We refer to these as *discordant triplets* and assume that in most cases the genetic regulation of the transcript and protein occur independently.

#### Discordant triplets

Applying three-choice model selection to the discordant triplets, we found that 99% were classified as Independent in the CC and 98% were classified as Independent in the DO (Table 3A). With the expanded model options, the Independent model was selected for 87% of triplets in both the CC and DO. The Complex model was selected for most of the remaining triplets in both studies. The high rate of correct classification of the discordant triplets indicates that there is little measurement error in the genotypes (middle variable) for both studies.

**Table 3.**
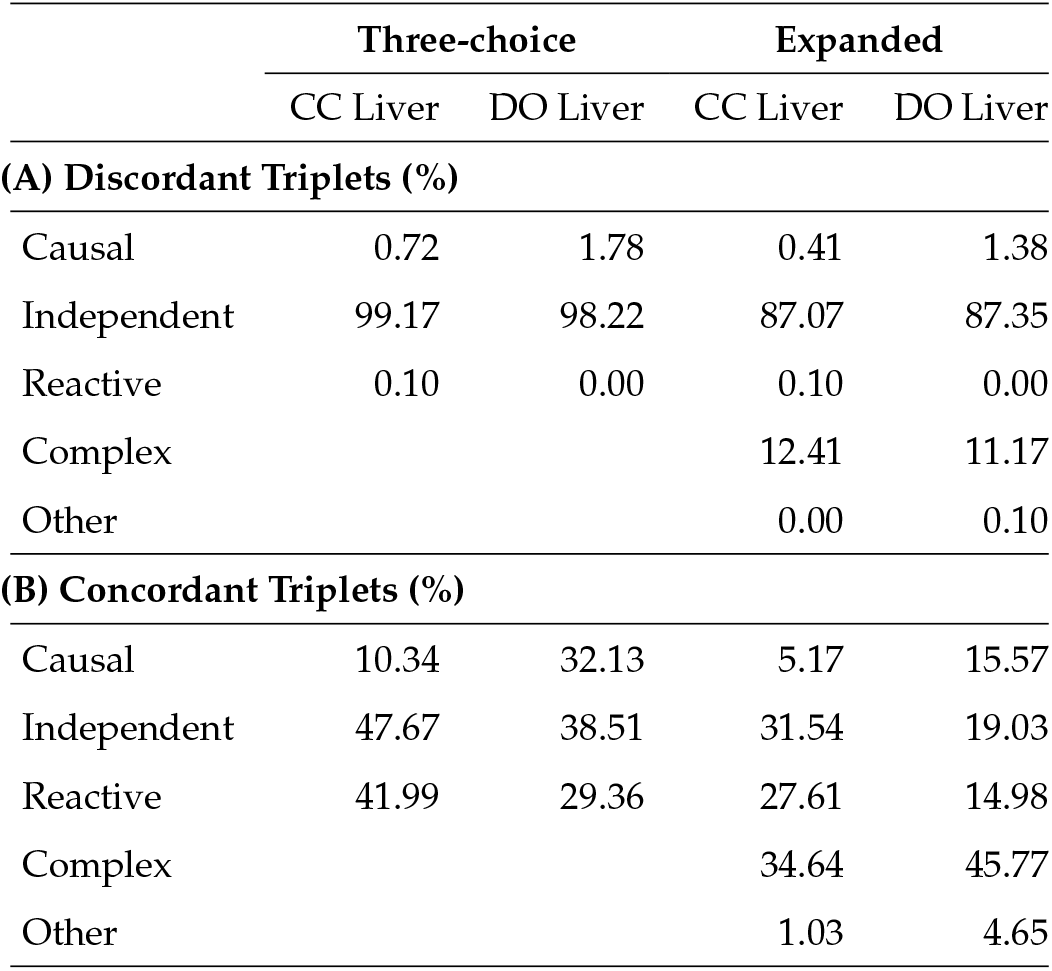
Classification of concordant and discordant triplets under three-choice and expanded model options.

#### Concordant triplets

Applying three-choice model selection to concordant triplets, we found that only 10% of triplets in the CC study and 32% of triplets in the DO study were classified as Causal (Table 3B). The remaining triplets were split between the Independent and Reactive models with a larger proportion of Independent classifications. Notably, more CC triplets were classified as Reactive compared to the DO triplets. With the expanded model options, the Causal model was selected for only 5% of CC triplets and 16% of DO triplets, and the Complex model was selected for 35% of CC triplets and 46% of DO triplets. Partial mediation, in which the QTL has direct effects on both the transcript and protein is possible. However, we expect that many of the triplets that were classified as Complex are due to un-modeled measurement error in the transcript data.

#### Allele Effects

The 8-state exogenous variable representing the DO genotypes provides more information than is available for univariate *X* (normal or binary). The relationships between the multi-state genotype and the univariate transcript and protein abundances are defined by eight regression coefficients (with mean zero constraint) that represent additive allele effects. If the causal structure is Independent, we expect the correlation between the two vectors of allele effects for *M* and *Y* to be randomly distributed, as was seen for the discordant triplets (Figure 7A). If the causal structure is Causal or Reactive, we expect the allele effects to be correlated as was seen for the concordant triplets (Figure 7B). However, nearby genes may have similar genetic effects that can result in correlation between the RNA and protein allele effects as we saw for discordant triplets that were classified as Causal (Figure 7A). Thus, while inconsistent allele effects can rule out a Causal relationship, consistent allele effects between Independent *M* and *Y* could be coincidental.

**Figure 7.**
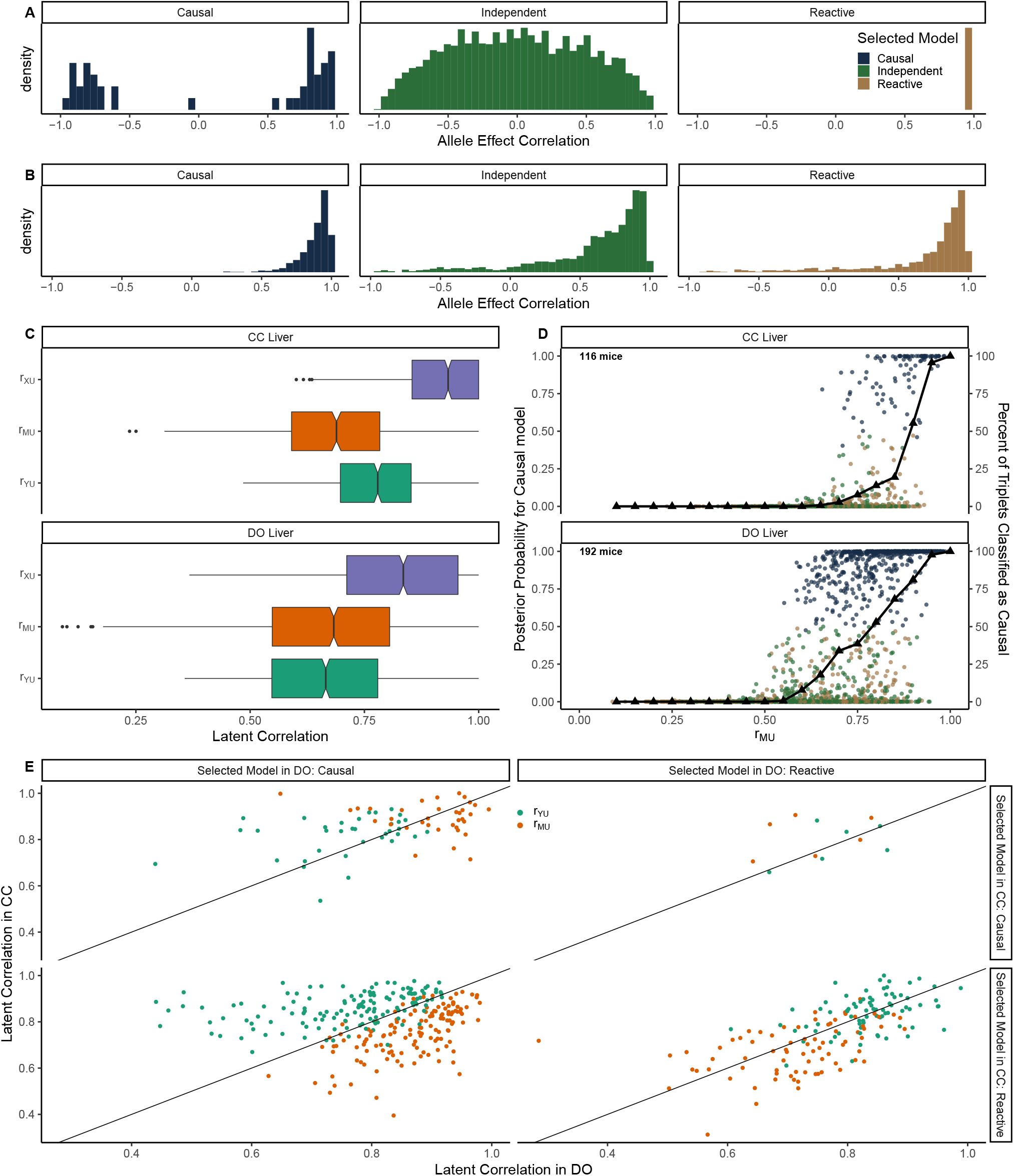
Mediation analysis of concordant and discordant triplets. Distribution of the correlation of eQTL and pQTL allele effects for discordant (A) and concordant (B) triplets, stratified by selected model. (C) Distribution of the estimated latent correlations in the CC (top) and DO (bottom) concordant triplets. (D) Results of three-choice Bayesian model selection for concordant triplets in the CC (top) and DO (bottom). Circles represent posterior probability for the Causal model as a function of the estimated latent correlation *r_MU_*, colored by selected model. Black triangles denote the proportion of triplets for which the Causal model was selected at values of *r_MU_* rounded to the nearest 0.05. (E) Comparison of estimated latent correlations *r_MU_* and *r_YU_* for concordant triplets present in both the DO and CC, classified as Causal or Reactive, and stratified by selected model in each population.

#### Diagnosis of mis-classified concordant triplets

We examined the estimated latent correlations of concordant triplets (Figure 7C). Assuming that the Causal model is true, *r_MU_* is an estimate of error in the transcript. The frequency of correct classification as a function of *r_MU_* was consistent with our simulated data, as can be seen by comparing Figure 5A with Figure 7D. As *r_MU_* increased, so did the proportion of triplets classified as Causal with three-choice model selection. The Causal model was not selected until *r_MU_* > 0.6, and when *r_MU_* > 0.9 almost 100% of triplets were classified as Causal. Thus, low measurement error in *M* supports correct inference of the Causal model with both univariate and multi-state *X*.

The misclassification of concordant triplets as Independent is consistent with low genotyping error in both studies (Figure 6B), but why are there more Reactive classifications in the CC than DO? To answer this question, we looked at 254 concordant triplets that were classified as either Reactive or Causal in both studies. Assuming the Causal model is true, greater error in the transcript data relative to protein data and a strong causal correlation between the transcript and protein data would result in *ρ_YU_* > *ρ_MU_* and a Reactive classification (Figure 6C). Recall that for the Causal model, *ρ_YU_* = *ρ_Y*Y_* · *ρ_Y*M*_* (Table 1). If we assume that the causal correlations between RNA and protein are similar between the CC and DO (Keele *et al*. 2021), differences in estimated *r_YU_* are largely due to measurement error in the proteins (*Y*). We saw that triplets classified as Causal in the DO and Reactive in the CC generally had a weaker *r_MU_* and stronger *r_YU_* in the CC than DO (Figure 7E, bottom left plot), consistent with more measurement error in the DO proteins relative to the CC. For the smaller number of triplets classified as Causal in the CC and Reactive in the DO, the reverse was true (Figure 7E, top right plot). When the selected model was the same in the CC and DO, the error correlations for *Y* and *M* were similar across the studies (Figure 7E, diagonal plots). For triplets classified as Causal, values of *r_MU_* were generally larger than values of *r_YU_*, and vice-versa for triplets classified as Reactive. This suggests that the higher rate of Reactive classifications in the CC is the due to less measurement error in the CC proteomics data. We note that the CC study used improved mass-spectrometer technology and replicated genotypes (Keele *et al*. 2021). Ironically, greater precision in protein measurement resulted in a higher rate of misclassification of concordant triplets in the CC data. A precisely measured *Y* and higher measurement error in *M* can result in incorrect Reactive classifications because *Y* is more strongly correlated with *M*\* than is *M* (Rockman 2008) (Figure 6C).

### Case Studies: Diagnosing mediation analysis

We selected three cases, two from mouse studies and one from a study of human cell lines, where genetic mediation analysis results appeared questionable. To evaluate each case, we determined ranges of the measurement error model parameters for each causal structure of interest that could have given rise to the observed data. To do this, we bootstrapped the data and computed a 95% support region for the data correlations (see Methods). We then selected simulated data sets that generated estimated data correlations within the support interval and noted the range of measurement error model parameters across the selected simulations. Lastly, we made a qualitative evaluation of the causal and error correlations considering the biological context and properties of the measurement technologies.

#### Mediation of distal eQTL in DO kidney

We obtained gene expression data from kidney tissue of 188 DO mice (Takemon *et al*. 2021) and examined a locus on chromosome 13 where a distal eQTL for *Sfi1* and a local eQTL for *Rsl1* co-map (Figure 8A, B). *Rsl1* is a transcription factor and a biologically plausible negative regulator of *Sfi1* (Krebs *et al*. 2012). We applied Bayesian model selection at the chromosome 13 locus (X) to evaluate *Rsl1* (M) as a candidate mediator of *Sfi1* expression (Y). We note that *Sfi1* also had a local eQTL on chromosome 11, which we included as a covariate along with age and sex in the Bayesian model selection. Three-choice model selection strongly favored the Independent model (posterior probability = 0.977), and expanded model selection favored the Complex model (posterior probability = 0.846), followed by the Independent model (posterior probability = 0.150) (Figure 8C). The Complex and Reactive models, if true, would require a direct causal effect of the chromosome 13 locus on the distal gene expression of Sfi1, so we focused on the Causal and Independent models.

**Figure 8.**
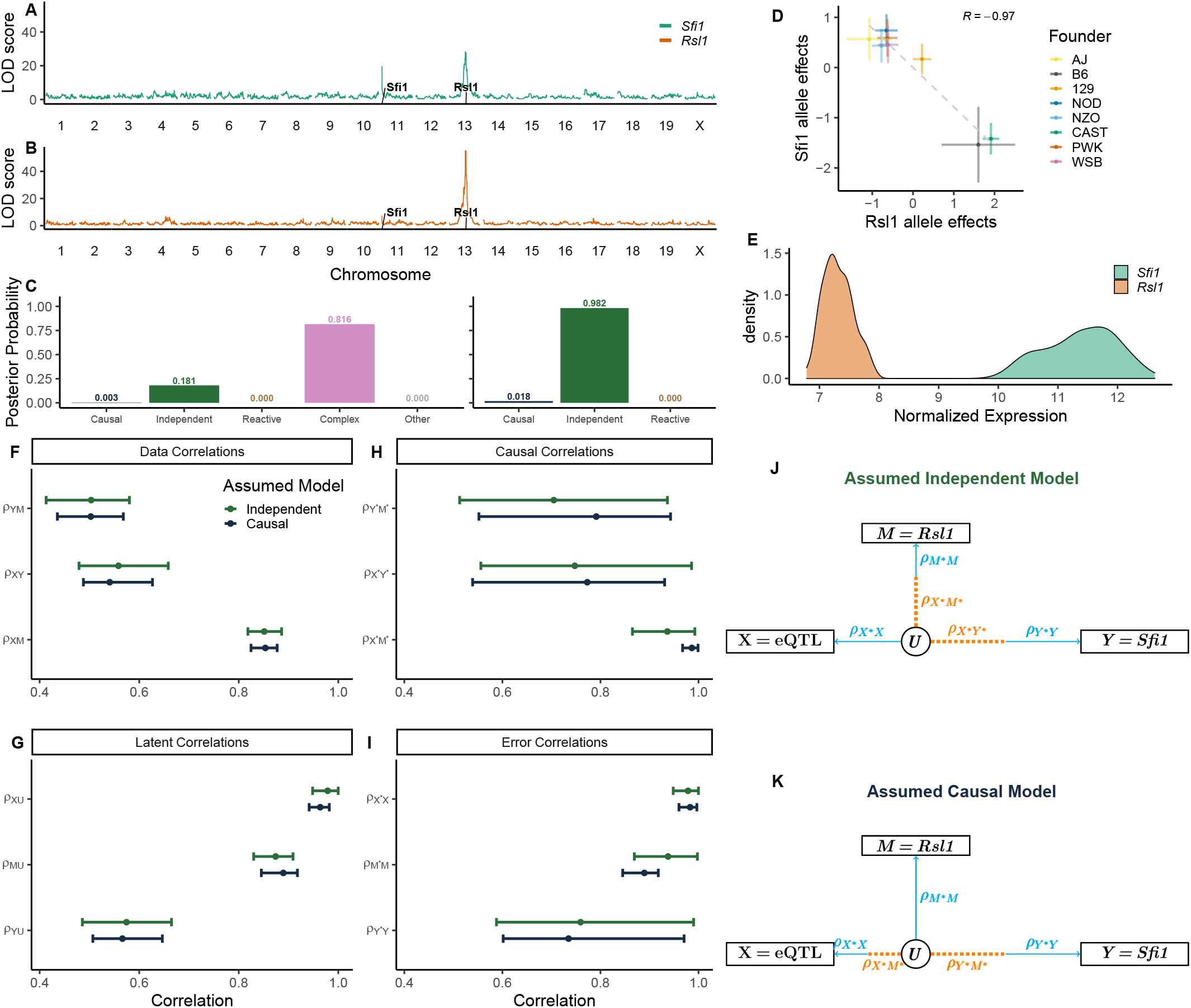
Mediation of *Sfi1* expression in DO kidney tissue. (A) *Sfi1* has a local eQTL on chromosome 11 and a distal eQTL on chromosome 13. (B) The distal eQTL co-localizes with a local eQTL for the transcription factor *Rsl1*. (C) Posterior model probabilities for the structure of the relationship between *Rsl1* and *Sfi1* calculated by Bayesian model selection with the expanded (left) and three-choice (right) model options. (D) Chromosome 13 QTL allele effects for *Rsl1* and Sfi1. (E) Variance stabilized transformed expression of *Rsl1* and Sfi1. (F-I) Median and 95% highest density interval for data, latent, causal, and error correlations corresponding to bootstrap estimated data correlations for Independent (green) or Causal (blue) model structures. DAGs of the latent variable model for median causal and error correlations assuming Independent (J) or Causal (K) model structure.

We bootstrapped the data and compared the measurement error parameters for the Independent and Causal models. The ranges are substantially overlapping, with the exception that *ρ_X*M*_* ≈ 1 under the Causal model (Figure 8F-I). If the Causal model is true, a combination of high measurement error on *Rsl1*, low genotyping error, and a strong causal genetic effect on *Rsl1*, such that *ρ_M*M_* < *ρ_X* X_* · *ρ_X*M*_*, would result in selection of the Independent model (Figure 8J, K). We note that *Rsl1* was expressed at low levels and may therefore have high measurement error; the *Rsl1* transcript also has a strong eQTL (LOD > 40), which is consistent with a strong causal correlation (*ρ_X* M*_*). The allele effects of the two transcripts were strongly anti-correlated (r = −0.96, p = 7.4e-05) (Figure 8D), consistent with a Causal model. We conclude that a Causal relationship in which *Rsl1* negatively regulates *Sfi1* is plausible and that selection of the Independent model was likely the result of measurement error in *Rsl1*.

#### Epigenetic mediation of gene expression in human lymphoblastoid cell lines

A study of epigenetic regulation of transcription in 63 human Lymphoblastoid Cell Lines identified genetic variants that affected gene expression (eSNPs) and chromatin accessibility (cSNPs), including a SNP in an interferon-stimulated response element (ISRE) in the first intron of *SLFN5* that is both an eSNP for *SLFN5* expression and a cSNP for a chromatin peak at the ISRE (Degner *et al*. 2012) (Figure 9). The position of the peak suggests that chromatin state mediates expression of *SLFN5*, which is an interferon-regulated gene, by controlling the accessibility of the ISRE to transcription factors (Mavrommatis *et al*. 2013). Bayesian model selection with three-choice model options selected the Reactive model (posterior probability 0.87), implying that the gene transcript mediates the local chromatin accessibility. Expanded model selection placed most of the posterior probability on the Complex model (0.82), followed by the Reactive model (0.156). The Reactive and Complex models cannot be ruled out, but the Causal model, in which the chromatin state mediates gene expression, has greater biological plausibility.

**Figure 9.**
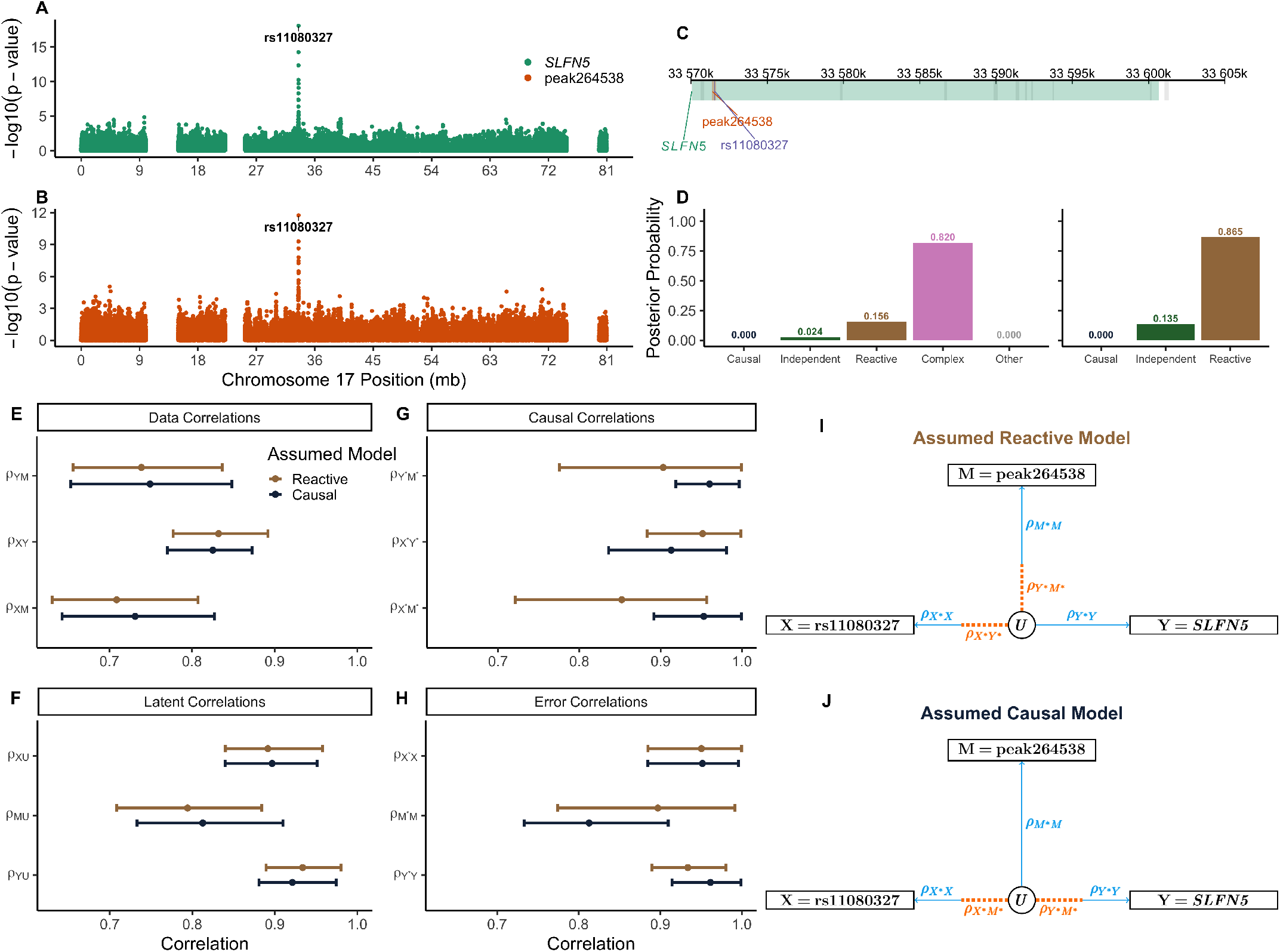
Mediation of *Slfn5* expression by a nearby chromatin peak in LCL data. rs11080327 is an eSNP for *SLFN5* (A), and a cSNP for a chromatin peak in the first intron (B). (C) The locations of *SLFN5*, rs11080327, and chromatin peak264538 on chromosome 17. (D) Posterior model probabilities for the relationship between the chromatin peak and *SLFN5* calculated by Bayesian model selection with the expanded (left) and three-choice (right) model options. (E-H) Median and 95% highest density interval for distributions of data, latent, causal, and error correlations for measurement error models that could generate the observed data when the causal structure is assumed to be Reactive (gold) or Causal (blue). DAGs of the latent variable model show relative strengths of causal and error correlations that could produce the data if the assumed model is Reactive (I) or Causal (J).

Bootstrapping the correlation parameters (Figure 9E-H) shows that under the Reactive model, the causal correlations must be weaker compared to the Causal model. For either model, measurement error in the SNP and in *SLFN5* expression is low and there is more measurement error in the chromatin peak. If the Causal model is true, there must be a strong causal correlation between the SNP and chromatin peak (*ρ_X* M*_*) and also between the chromatin peak and the target *SLFN5* (*ρ_Y*M*_*) (Figure 9J). This is consistent with the strong genetic associations for both the chromatin peak and *SLFN5* (— log_10_ (*p*–value) > 10). Alternatively, if the Reactive model is true, the causal correlations *ρ_X* M*_* and *ρ_Y*M*_* must be weak, and there must be less error in the chromatin peak and more error in *SLFN5* compared to the Causal model. In light of our expectation that chromatin data are noisy and that open chromatin regulates transcript abundance, we suspect that the mediation analysis inference of a Reactive relationship is incorrect and that the true causal structure is Causal or possibly Complex.

#### Mediation of distal pQTL in DO Liver

In the DO liver data (Chick *et al*. 2016) we looked at protein to protein mediation and observed a distal pQTL for TUBG1 that co-maps on chromosome 8 with a local pQTL for NAXD (Figure 10). The allele effects for TUBG1 were negatively correlated with those of NAXD (r = −0.89, p = 0.003). We applied Bayesian model selection to determine if NAXD could be a mediator of the TUBG1 QTL and confirmed that the greatest posterior probability was on the Causal model for both three-choice (0.978) and expanded model options (0.694). Bootstrapping showed overlap between the distribution for *ρ_XY_* and *ρ_YM_* as the weakest data correlation, indicating the data may be consistent with either the Causal or Independent model. Examination of the error model parameter ranges indicated that, if NAXD and TUBG1 are Independent (Figure 10L), there must be a strong causal correlation between the QTL and NAXD (*ρ_X* M*_*), little error in the genotype, and little error in NAXD. Alternatively, if NAXD is a mediator of the TUBG1 QTL (Causal model, Figure 10M), the strength of the causal correlation between the QTL and NAXD must be even stronger, with slightly less genotyping error, but more error in NAXD.

**Figure 10.**
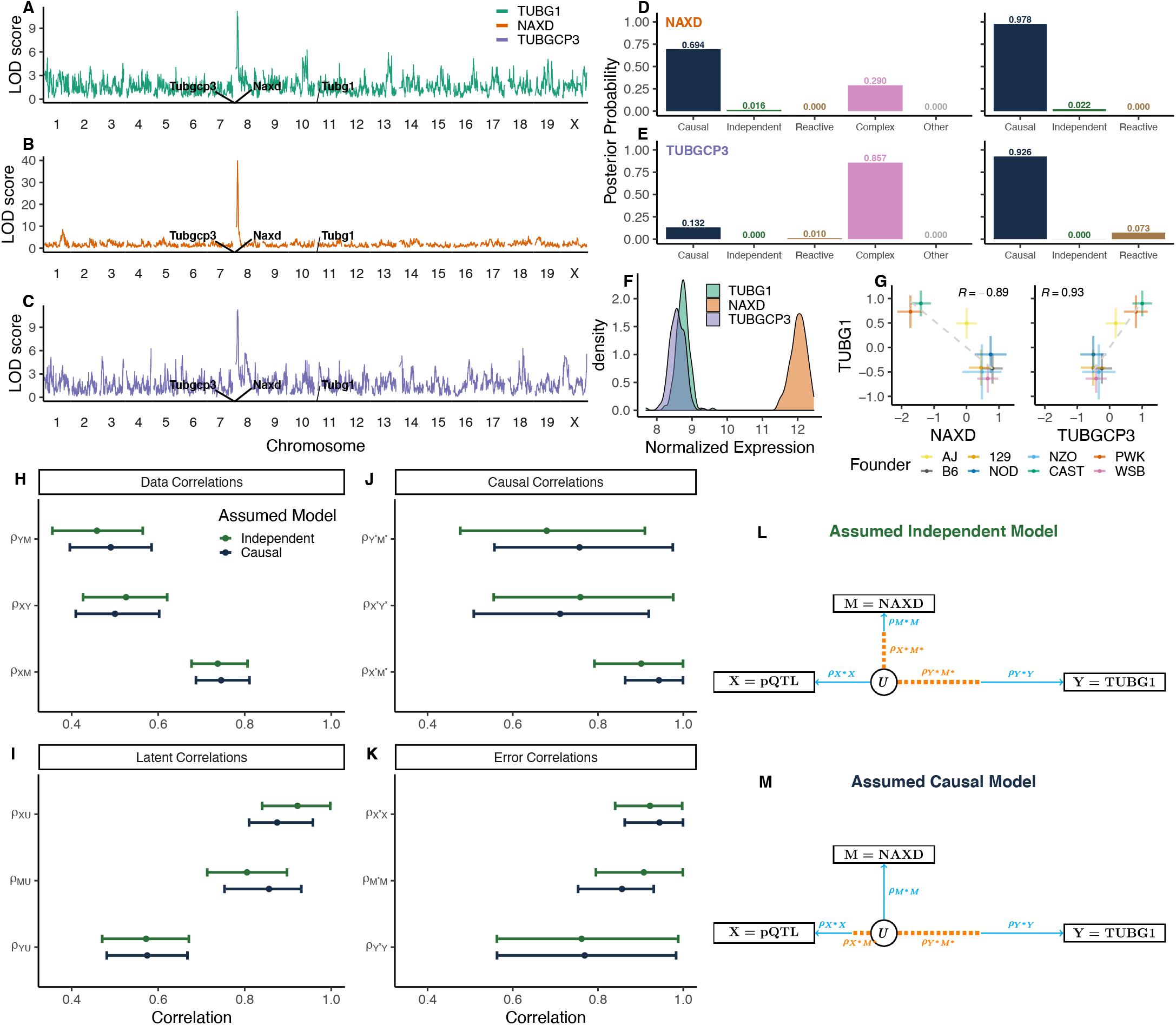
Mediation of TUBG1 protein abundance in DO liver tissue. A distal pQTL for TUBG1 (A) co-localizes with chromosome 8 local pQTLs for NAXD (B) and TUBGCP3 (C). Posterior probabilities for NAXD (D) and TUBGCP3 (E) as candidate mediators of TUBG1, calculated by Bayesian model selection with expanded (left) and three-choice (right) model options. (F) Normalized protein abundance of NAXD, TUBGCP3, and TUBG1. (G) Allele effects for TUBG1 compared to those for NAXD (left) and TUBGCP3 (right). (H-K) Median and 95% highest density interval for distributions of data, latent, causal, and error correlations for measurement error models that could generate the observed data when the causal structure is assumed to be Independent (green) or Causal (blue). DAGs of the latent variable model show relative strengths of causal and error correlations that could produce the data if the assumed model is Independent (L) or Causal (M).

At this point, we might have concluded that NAXD is a mediator. However, the chromosome 8 pQTL for TUBG1 also co-maps with a local pQTL for TUBGCP3 and the allele effects of the pQTL are positively correlated (r = 0.93, p = 0.0008). TUBGCP3 and TUBG1, together with a third protein TUBGCP2, form the *γ*-tubulin small complex (Oakley *et al*. 2015; Farache *et al*. 2018). This functional relationship suggests that TUBGCP3 likely mediates the distal pQTL for TUBG1. Three-choice Bayesian model selection to test TUBGCP3 as a mediator of the TUBG1 pQTL showed that Causal model had the greatest posterior probability (0.926). Under the expanded model options, the Complex model was selected (posterior probability 0.857). In addition, analysis of CC mice (Keele *et al*. 2021) supported TUBGCP3 as a Causal mediator.

Considering the biological evidence supporting TUBGCP3 as a mediator of TUBG1, why does mediation analysis provide stronger support for NAXD? The NAXD QTL (LOD ≈ 40) was much stronger than that of TUBG1 (LOD ≈ 13). In addition, NAXD was more abundant than the target gene TUBG1 (Figure 10F), indicating that TUBG1 may be measured with more error. The allele effects of the Chr 8 pQTL on NAXD and TUBGCP3 are highly similar so we cannot rule out either candidate based on allele effects. The nearly 100% posterior probability for mediation by NAXD creates misleading certainty but it is based on a model that does not account for measurement error. This example illustrates a common scenario in which mediation analysis supports a highly expressed gene with a strong local QTL as a mediator when the true relationship is Independent.

## Discussion

In this work, we examined the impact of measurement error on mediation analysis. We showed that in the presence of measurement error, data from any three variable causal structure will be consistent with partial mediation, *i.e*., the Complex model with no measurement error. It follows that mediation analysis, which does not account for measurement error, can infer partial mediation even in the absence of an indirect effect of *X* on *Y* through *M*. This outcome becomes more likely as sample size increases, which is especially concerning for methods such as the Sobel test that focus solely on detecting the indirect effect.

The measurement error model with three observable variables is not identifiable, *i.e*., it is not possible to uniquely estimate each of the model parameters. We introduce a latent variable model with estimable parameters and illustrate how they relate to the causal and error correlation parameters of the measurement error model. Using the latent variable model, we identify scenarios for which a three-choice model selection strategy (excluding partial mediation) can lead to consistent or inconsistent identification of causal structure. Measurement error in the middle variable is most critical, but there are many other scenarios that can result in inconsistent inference (Table S1).

Genetic mediation analysis has some unique features that distinguish it from more general applications of mediation analysis. The candidate mediator is typically a transcript or protein whose coding gene co-locates with the target QTL. If the target is a molecular trait from a distant gene, it seems reasonable that any causal effect linking the mediator to the target would be from *M* to *Y*, and the Reactive model would be *a priori* unlikely. On the other hand, if the target is a clinical trait or a molecular trait that co-locates with the mediator, we cannot rule out the Reactive model. When mediation analysis unexpectedly indicates the Reactive model, it is likely due to low measurement error in the target as illustrated by our analysis of concordant triplets. Multi-state genotypes, as in our DO and CC mouse examples, can reduce the rate of misclassification by providing information that is not available with bi-allelic variants. We have previously shown, using simulations with no measurement error, that multi-state genotypes can reduce false detection of mediation and provide stronger evidence of mediation when it is present (Crouse *et al*. 2022). In the presence of measurement error, when the QTL allele effects for a candidate mediator and a target do not align, we can confidently rule out mediation. However, as was shown in our third case study, matching allele effects can be misleading.

Our case studies illustrate the most common failure modes of genetic mediation analysis. In the first case study, a likely Causal relationship is classified as Independent (Figure 6B). Correlated allele effects provide a clue that the Independent classification may be in error and experimental evidence from other studies indicates that the transcript factor *Rsl1* regulates downstream gene expression (Krebs *et al*. 2012). In the second case study, a likely Causal relationship is classified as Reactive (Figure 6C). The coding gene for *SLFN5* and the chromatin peak at the ISRE co-locate with the target QTL, so we cannot rule out the Reactive model. Our conclusion that the true relationship is Causal relies on our assumptions that chromatin data are noisy and that chromatin states regulate gene expression and not vice-versa. In the third case study, an Independent relationship is classified as Causal (Figure S5B). We were able to rule out NAXD as a mediator because there is another candidate mediator (TUBGCP3) with greater biological plausibility. Recall that in our analysis of discordant triplets, mediation analysis correctly classified >98% of the triplets as Independent. The difference here is that NAXD was not chosen at random; it was identified by testing each of the genes in a QTL support interval to identify the protein with the strongest evidence for Causal mediation. This afforded multiple opportunities to encounter a candidate mediator with the same allele effects as the true mediator. If this study had relied on binary genotypes, any nearby gene with a strong local pQTL could be mistakenly identified as a Causal mediator. In each case study, we were able to diagnose the problem by considering how measurement error could have led to selection of an incorrect causal structure. However, the key to spotting these problems was the existence of independent evidence or prior knowledge about the biological mechanism of regulation that was at odds with the mediation analysis inference.

The ideal solution to these challenges would be to minimize measurement error. It is also possible, at additional cost, to design experiments with technical replication to estimate measurement error. Aygün *et al*. (2022) used technical replicates in a gene expression study to estimate measurement error in the mediator and target. They excluded triplets with unbalanced error from mediation inference, which reduced the rate of false Reactive classifications. Methods to obtain direct estimates of measurement error have been proposed for proteomics data (Peshkin *et al*. 2019), and it seems likely that related approaches could be developed to estimate the precision of RNA-Seq data. Incorporating estimates of measurement error could enable accurate estimation of causal correlations.

We limited the scope of our mediation analysis to only three variables and to univariate measures, with the exception of multi-state genotypes. Expanding the causal system to include more variables or using multivariate observations in mediation analysis could mitigate some of the concerns raised here. For example, bidirectional mediation analysis (Talluri and Shete 2018) of the relationship between *Rsl1* and *Sfi1* including the QTL on chromosome 11 (local to Sfi1) should correctly identify the causal structure. In addition, we made simplifying assumptions including independent measurement error and the absence of hidden confounders.

The problem of hidden confounders of *M → Y* has received a great deal of attention (Saunders and Blume 2018). Mendelian randomization (MR) (Katan 1986; Didelez and Sheehan 2007; Richmond *et al*. 2016) is a form of causal inference that is robust to the presence of hidden confounders, but it achieves robustness by assuming that there is no direct effect between *X* and *Y*. Thus MR cannot distinguish between the Independent and Causal mediation models (Crouse *et al*. 2022). For example, MR would fail to distinguish among candidate mediators with correlated allele effects, likely favoring the one with the strongest local QTL because any information *M* possesses for *Y* after accounting for *X* is discarded. While MR does not address our objectives, it is a powerful and widely used method of causal inference and the potential impact of measurement error on MR warrants further study.

Genetic mediation analysis is an effective approach to identify candidate genes and to generate hypotheses about the mechanism underlying the effects of genetic variation. While the pitfalls of mediation analysis are real, we hope that this examination of the impact of measurement error will support more informed application, acknowledging weaknesses, while not detracting from its utility. As a simple approach to evaluate a mediation analysis inference, we recommend transforming the estimated data correlations to latent correlations and then, assuming each causal structure in turn, consider whether the relative sizes of the causal and error correlations are consistent with prior knowledge about the biological system and measurement technologies. Bootstrapping can be helpful for investigating how precise the estimated data correlations are. As illustrated in our case studies, it may not be possible to confidently establish (or rule out) mediation based solely on three-variable data. Incorporating more variables or employing experimental designs that support the estimation of measurement error could mitigate some of these challenges. However, in the absence of additional evidence to support a mediation inference, there may be no substitute for independent experimental validation.

## Methods

### Simulations

We simulated data from the measurement error model with a univariate exogenous variable that was normally distributed. For each of the Causal, Independent, and Reactive models, 100,000 configurations were constructed by randomly sampling two causal and three error correlations from a beta distribution: *ρ ~* Beta (5,1.25). We chose this as a realistic distribution for causal and error correlations with mean 0.8 and 95% highest density interval between 0.5 and 1, allowing for weak correlations but placing greater density on moderate to strong correlations. We compared the distributions of data correlations to those observed in the DO liver data to confirm biological plausibility. Then *X*\* was sampled from a standard normal distribution and *M\** and *Y\** were simulated according to the causal correlations using the method described in (Crouse *et al*. 2022) with the effect size calculated as *ρ*^2^. We used the same method to simulate *X*, *M*, and *Y* from their causal counterparts with the desired error correlation.

### Mediation Analysis with Bayesian Model Selection

We used Bayesian model selection implemented in the bmediatR R package (Crouse *et al*. 2022) to obtain the posterior probability for the standard mediation model (no measurement error) using either the three-choice or expanded model options. We ran bmediatR with default priors for the effect sizes, and a uniform prior over models. Data were classified according to the model with the greatest posterior probability. With the three-choice model options, the posterior probability was calculated for only the Causal, Independent, and Reactive models. For the expanded model options, the posterior probability was calculated for all models excluding models with an edge *Y* → *M* that are likelihood equivalent to models with *M* → *Y*. If the greatest posterior probability was not assigned to one of the Causal, Reactive, Independent, or Complex models, the data were classified as non-mediation.

### QTL Mapping Analysis

QTL mapping and allele effect estimation in mouse data were done using the qtl2 R package (Broman *et al*. 2019), which fits a linear mixed effect model that accounts for population structure encoded in a genetic relationship matrix, *i.e*., kinship matrix (Kang *et al*. 2010). Allele effects were estimated as best linear unbiased predictors (BLUPs) to stabilize estimates. Sex, diet, and litter were used as covariates in the DO liver data; sex was used as a covariate in the DO kidney data; and sex was used as a covariate in the CC liver data. QTL in the LCL data were identified by calculating the – log_10_ (*ρ – value*) from regressing chromatin data or gene expression onto the genotype of each SNP compared to a null model with no genotype term.

### Using bootstrap sampling to compare real data to simulated data

To identify measurement error model parameters that are consistent with the observed data, we used the boot R package (Davison and Hinkley 1997; Canty and Ripley 2021) to sample with replacement (10,000 times) to generate empirical distributions for the estimated data correlation matrix. If *X* was multivariate we used canonical correlation to estimate *ρ_XM_* and *ρ_XY_*. Then, we filtered the previously described measurement error model configurations used in data simulations to those that produced data correlations jointly within the range of the bootstrapped distribution.

## Data availability

All analyses were performed using the version 4.2.0 of the R statistical programming language (R Core Team 2022). All data and R code used to generate the results are available at figshare (https://doi.org/10.6084/m9.figshare.20126543).

Data are also available for download and interactive analysis with the QTLViewer webtool (Vincent *et al*. 2022) (https://github.com/churchill-lab/qtlapi) for the DO Liver (https://churchilllab.jax.org/qtlviewer/svenson/DOHFD) and DO Kidney (https://churchilllab.jax.org/qtlviewer/JAC/DOKidney) studies. The individual CC liver data are available in QTLViewer format from figshare (https://doi.org/10.6084/m9.figshare.12818717) at data/qtlviewers/cc_individuals_proteomics_qtlviewer.Rdata. Both genotype (Li *et al*. 2016) and RNA-seq data (Pickrell *et al*. 2010; van de Geijn *et al*. 2015) for Yoruba LCLs are publicly available for download (http://eqtl.uchicago.edu/jointLCL/). The DNase-seq data for the 69 cell lines with RNA-seq data were used as previously processed (Grubert *et al*. 2015).

## Appendix

### Violation of conditional independence between measured variables

Conditional independence between measured variables can be violated even when the underlying causal variables are conditionally independent. Suppose the causal structure is Causal such that the causal correlations satisfy the conditional independence constraint *ρ_X*Y*_* = *ρ_X*M*_* · *ρ_Y*M*_*, and assume that all of the causal and error correlations are non-zero. Conditional independence between *X* and *Y* hold if *ρ_XM_* · *ρ_YM_* = *ρ_XY_* and this implies that there is no measurement error in *M*, as

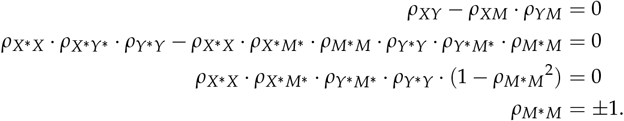

Similar algebra shows that if *ρ_M*M_* = ±1 and the structure of the causal system is Causal then *ρ_XY_* – *ρ_XM_* · *ρ_YM_* = 0. Thus, when the structure of the causal system is Causal, conditional independence between measured variables is satisfied if and only if *ρ_M*M_* = ±1, *i.e*., there is no measurement error in *M*. Repeating the above with the constraints for the Independent and Reactive models, we see that measurement error in the middle variable leads to lack of conditional independence between measured variables.

This result is overlooked in mediation analyses that rely on conditional independence criteria to infer complete mediation. For example, Chen *et al*. (2007) introduce the *Causality Equivalence Theorem* in which they prove that the causal relationship *X* → *M* → *Y* exists and there are no unmeasured confounders of *M* → *Y* if and only if the following three conditions hold: *X* → *Y*, *X* → *M*, and *X* ⊥ *Y*|*M* (conditional independence). The proof of this theorem assumes that all direct causes of each variable are measured without error. However, it is reasonable to assume that measurement error is present in any real setting and conditional independence between the exogenous variable and target will not hold in the measured data even if the causal structure is *X* → *M* → *Y*.

### Deriving latent correlations from the measurement error model

In our measurement error models, the data correlations can be described in terms of causal and error correlations (Equation 1) or in terms of latent correlations (Equation 2). These equivalencies can be used to express the latent correlations in terms of the causal and error correlations as follows,

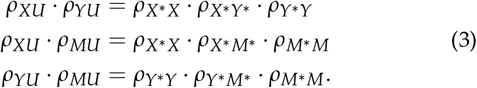

Solving this system for the latent correlations yields,

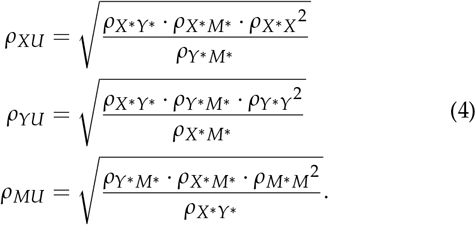

For the Causal model, *ρ_X*Y*_* = *ρ_X*M*_* · *ρ_Y*M*_* and Equation 4 simplifies to,

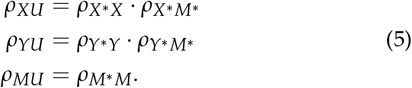

Table 1 summarizes the latent correlations for the Independent and Reactive models. In each case, the latent correlation for the middle variable is equivalent to the error correlation on that variable while the other latent correlations are products of one causal and one error correlation.

### The measurement error model likelihood

The log likelihood of the data given a model correlation matrix, Σ, is described by

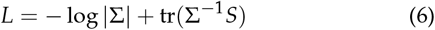

where *S*_3*x*3_ is the correlation matrix of the observed data. For the Casual measurement error model,

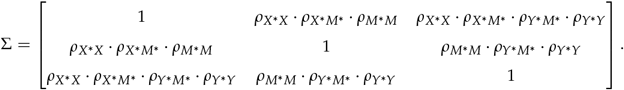

Each combination of measurement error model parameters generates one Σ, but the same Σ may be achieved by different parameter combinations. For example, every entry with *ρ_X* X_* includes the product *ρ_X* X_* · *ρ_X*M*_*. Thus, the values of *ρ_X*_* and *ρ_X* M*_* may be swapped, and the resultant Σ will not be changed. This property holds when the causal structure is Independent or Reactive, indicating that the measurement error model is unidentifiable and the causal and error correlations cannot be uniquely estimated from the data.

However, the latent variable model may also be used to describe the data and thus we can write Σ for any causal structure as follows,

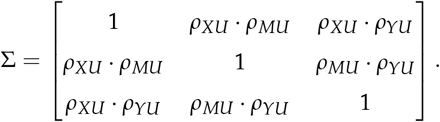

This formulation of Σ is identifiable and can be used to estimate the latent correlations via maximum likelihood estimation.

We numerically optimized the likelihood in Equation 6 using the “L-BFGS-B” method (Byrd *et al*. 1995) with bounds of (−1, 1) and initial condition 0.5. In the case when *X* is a multi-state variable, we estimate *r_XY_* and *r_XM_* with canonical correlations. Doing so results in underestimation of *ρ_XU_*, providing an upper bound on the amount of error in *X* (Figure S5).

### Bootstrapping procedure

We bootstrapped the data to obtain an approximate confidence region for the data correlation parameters. We then used simulations to determine the ranges of the measurement error model parameters that can produce data correlations falling within this confidence region. The procedure is as follows:

1. Sample the data with replacement 10,000 times.
2. Regress out any covariates used for mediation analysis.
3. Estimate the data correlations for each sample, using canonical correlation to estimate *r_XM_* and *r_XY_* if *X* is multivariate.
4. Define an elliptical region that captures 95% of absolute value of all boostrapped data correlations.
5. Filter the measurement error model simulations to find the parameters (causal and error correlations) that produce estimated data correlations within the bootstrap confidence region.
6. For each causal structure, compute the 95% highest density interval for each causal and error correlations over the filtered set of measurement error model simulations.

## Supplemental Materials

**Figure S1.**
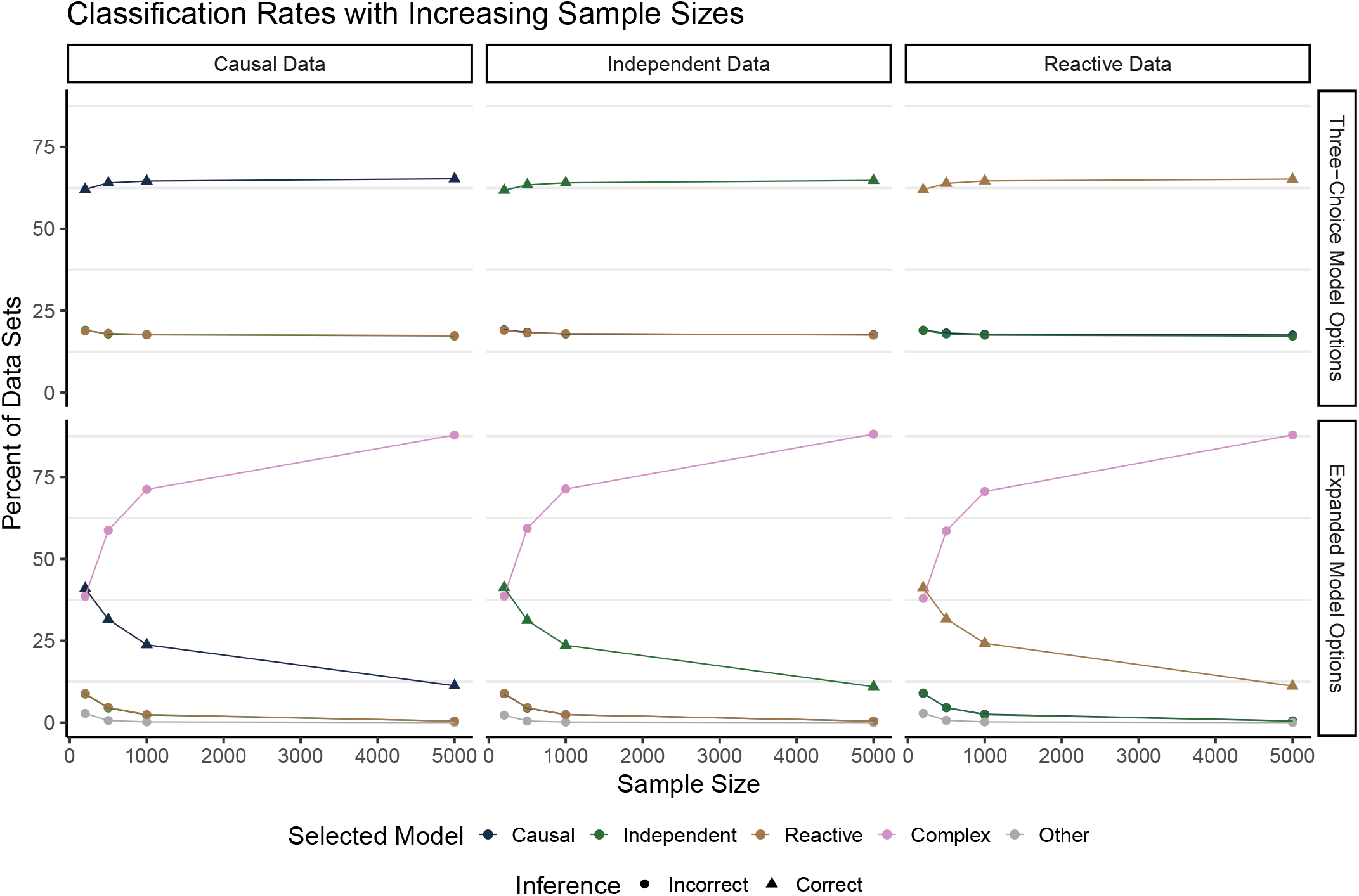
Classifications of simulated data with increasing sample sizes. Percent of data sets assigbned to each of the three causal structures (colored line) is shown as a function of sample size. Columns correspond to the simulated causal structure. Classification rates are shown for three-choice (top row) and expanded (bottom row) model options. The correct model is denoted with triangles and the incorrect models are denoted with circles.

**Figure S2.**
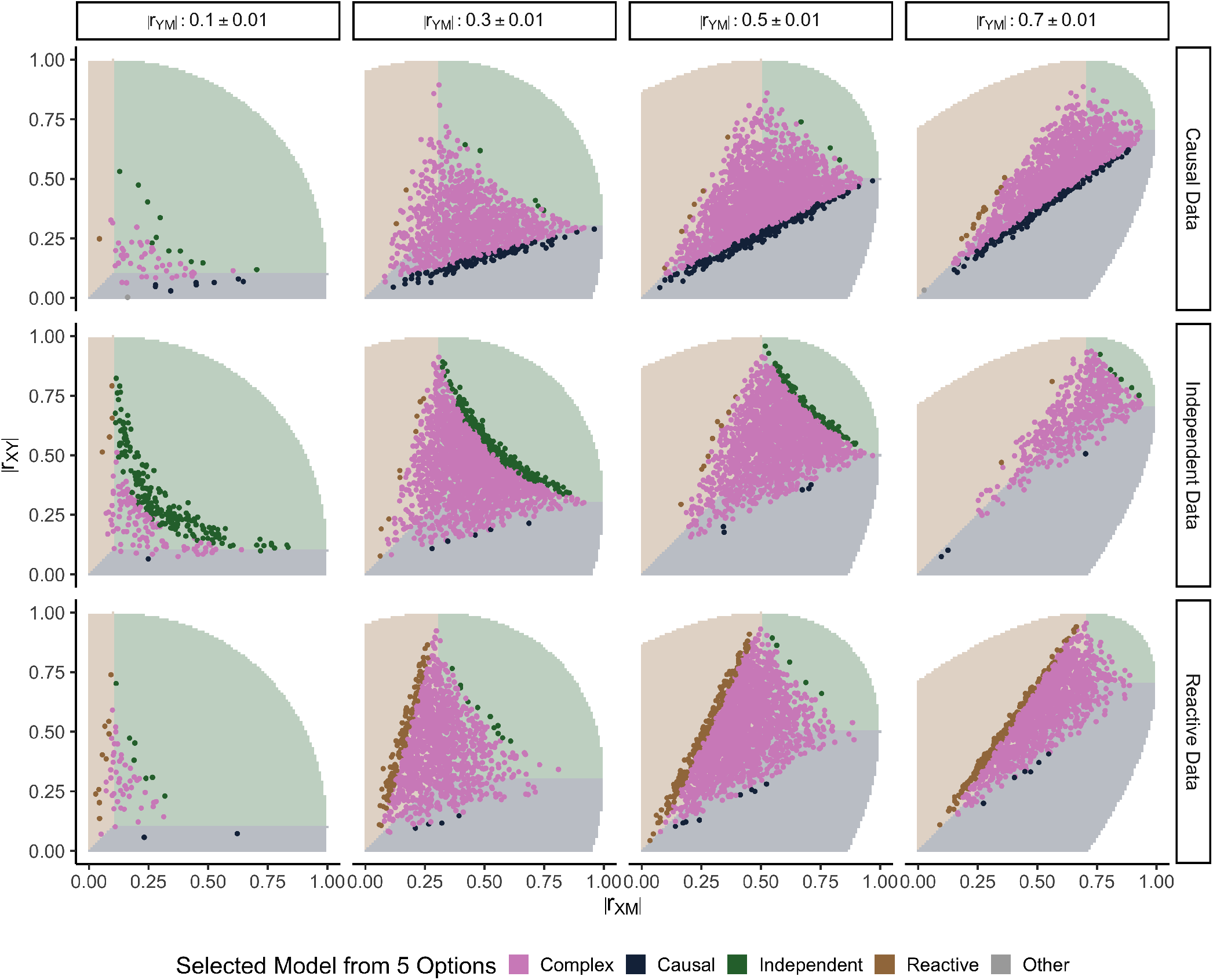
Estimated data correlations from simulations with *N* = 5000 and expanded model options. Each row of panels corresponds to simulations of a causal structure. Columns correspond to binned values of *r_YM_*. The x- and y-axes show *r_XM_* and *r_XY_*, respectively. Points representing the estimated data correlations are colored to indicate the model with the greatest posterior probability. Shaded regions indicate the three-choice model selection inference rule, and the unshaded region delineates where the correlation matrices are not positive semi-definite. See Figure 4 for results from simulations with *N* = 200.

**Figure S3.**
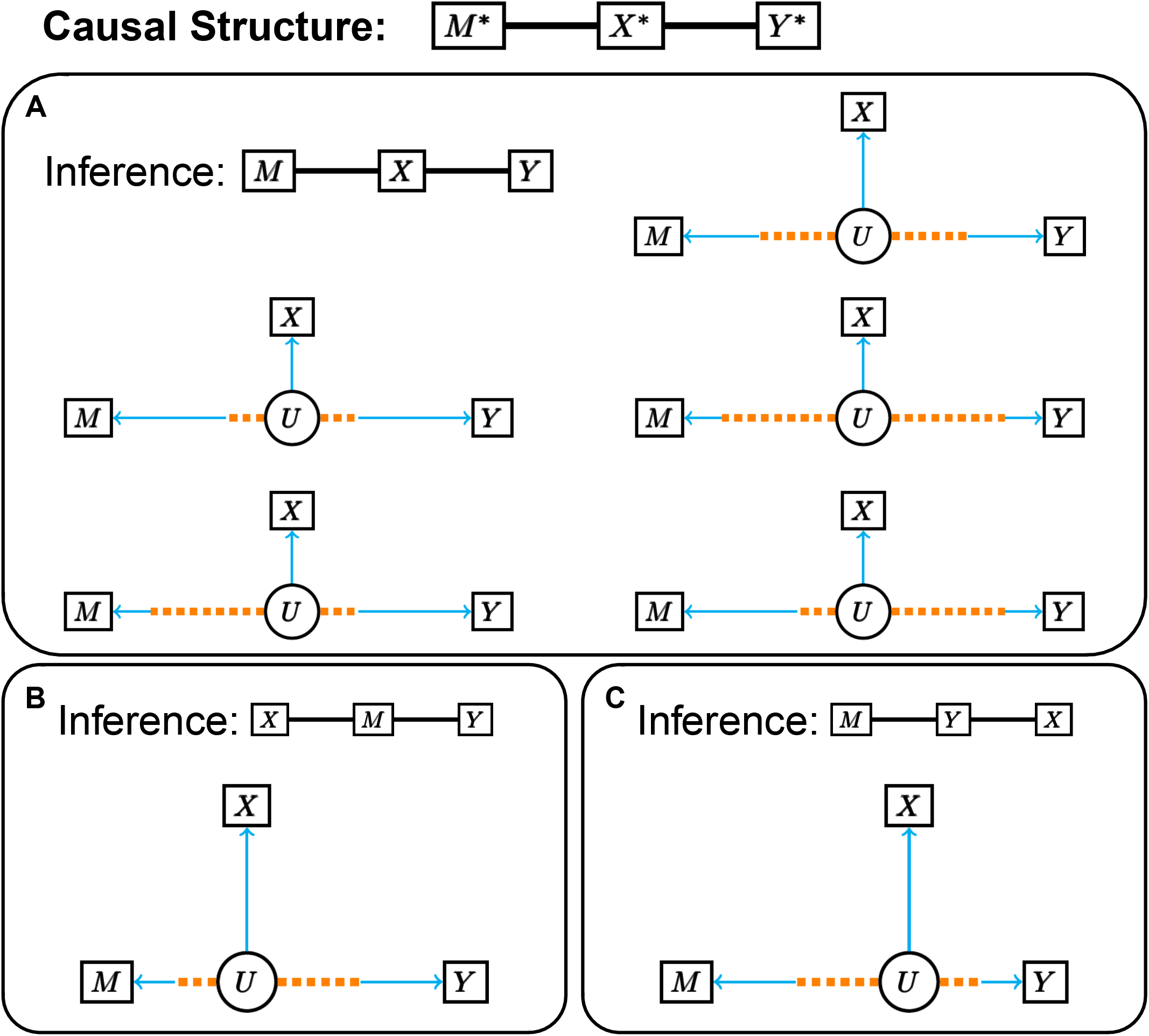
Latent variable representation of the Independent measurement error model. Configurations of the latent variable model representing the Independent model that result in (A) consistent and (B-C) inconsistent inferences. Figure designed the same way as Fig 6. (A) The correct model is inferred if the latent correlation *ρ_XU_* is the strongest (the latent variable arm for *X* is the shortest). This can be achieved if there is an equal amount of error in all three variables (top right) or if there is less error in *X* than *M* and *Y* (middle left). If *X* is noisier than *M* and *Y*, the correct model may still be inferred if the causal correlations are weak (middle right). The bottom row shows scenarios where *M* and *Y* satisfy different configurations. (B) The Causal model is inferred if *ρ_MU_* is the strongest. When the causal structure is the Independent model, this will only occur if *ρ_XU_* is weaker than both the error correlation and causal correlation contributing to *ρ_MU_*. The relative magnitude of the causal and error components of *ρ_MU_* do not matter as long as their product results in the strongest latent correlation. Similarly, the relative magnitude of the causal and error components of *ρ_YU_* do not matter as long as *ρ_YU_* is not larger than *ρ_MU_*. (C) Shows the analogous scenario to (B) for inferring the Reactive model by swapping *M* and *Y*.

**Figure S4.**
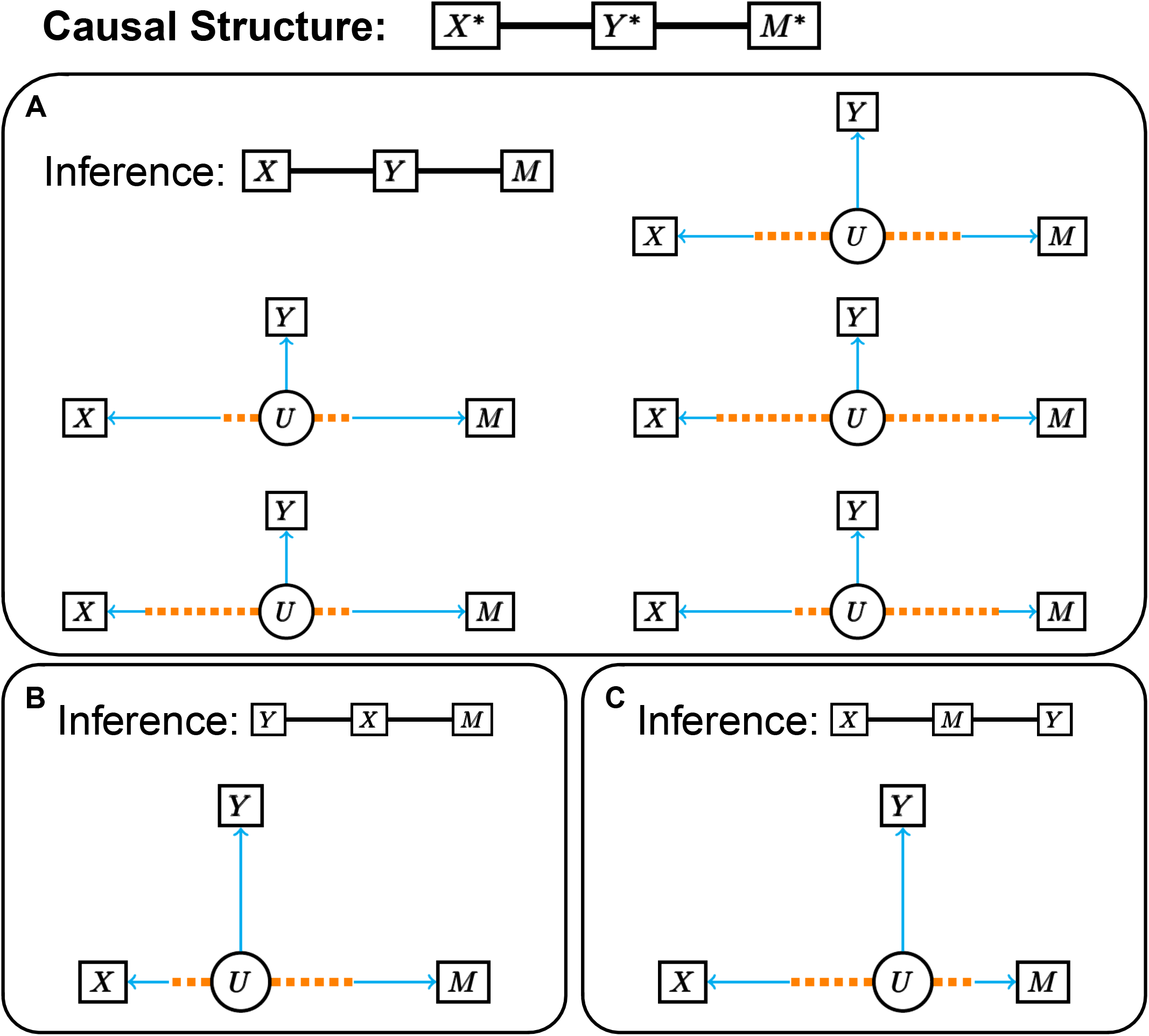
Latent variable representation of the Reactive measurement error model. Configurations of the latent variable model representing the Reactive model that result in (A) consistent and (B-C) inconsistent inferences. Figure designed the same way as Fig 6. (A) The correct model is inferred if the latent correlation for *Y* is the strongest (the latent variable arm for *Y* is the shortest). This can be achieved if there is an equal amount of error in all three variables (top right) or if there is less error in *Y* than *X* and *M* (middle left). If *Y* is noisier than *X* and *M*, the correct model may still be inferred if the causal correlations are weak (middle right). The bottom row shows scenarios where *X* and *M* satisfy different configurations. (B) The Independent model is inferred if *ρ_XU_* is strongest. When the causal structure is the Reactive model, this will only occur if *ρ_YU_* is weaker than both the error correlation and causal correlation contributing to *ρ_XU_*. The relative magnitude of the causal and error components of *ρ_X_* do not matter as long as their product results in the strongest latent correlation. Similarly, the relative magnitude of the causal and error components of *ρ_MU_* do not matter as long as *ρ_MU_* is not larger than *ρ_XU_*. (C) Shows the analogous scenario to (B) for inferring the Causal model by swapping *X* and *M*.

**Figure S5.**
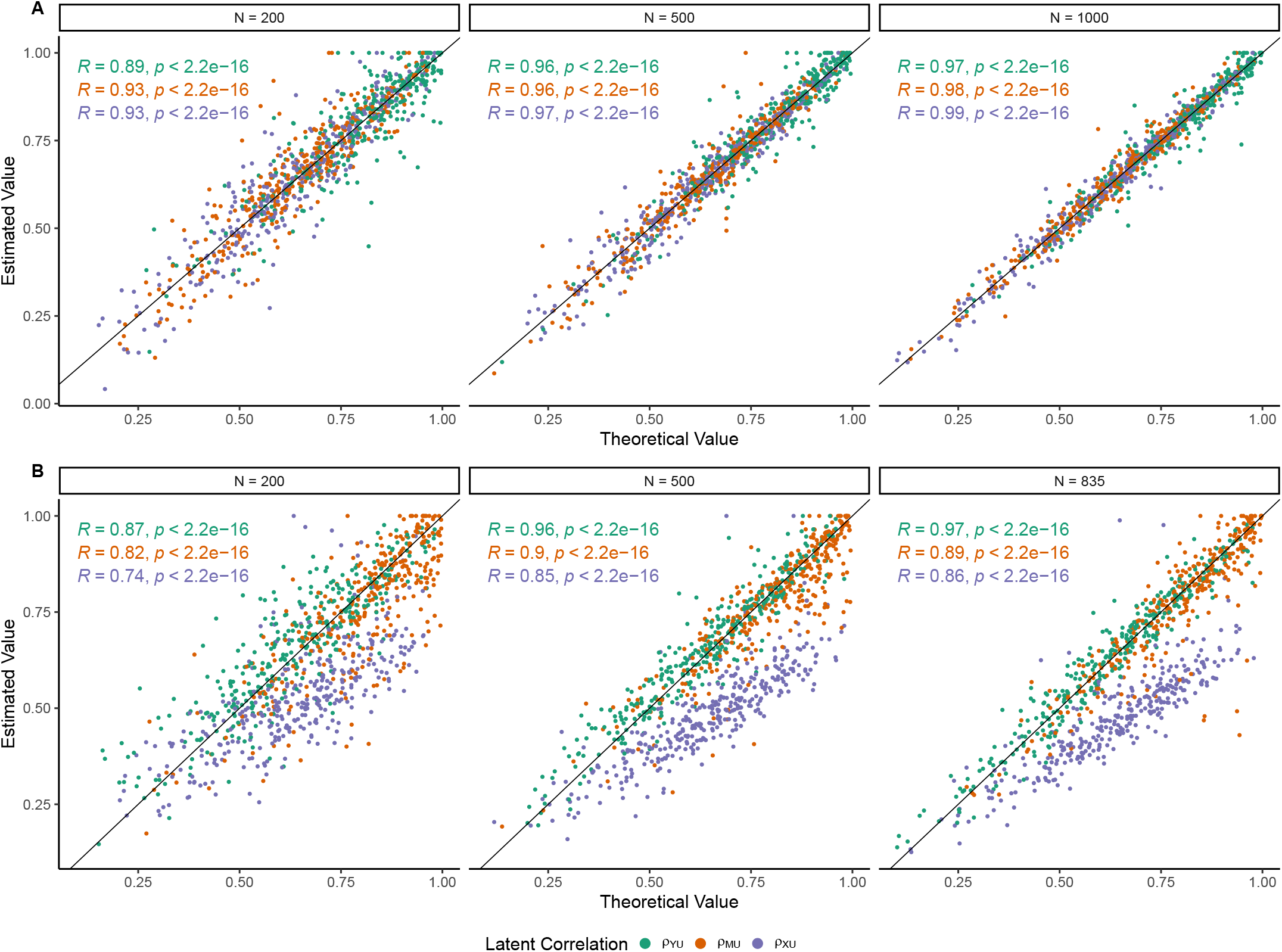
Latent correlations estimated from observed data. (A) Estimated latent correlations for a subset of 1000 simulations of the Causal measurement error model with univariate *X*. (B) Estimated latent correlations for simulations of the same measurement error models using multi-state *X* randomly selected from the genotype probabilites from liver tissue of 835 DO mice.

**Table S1.**
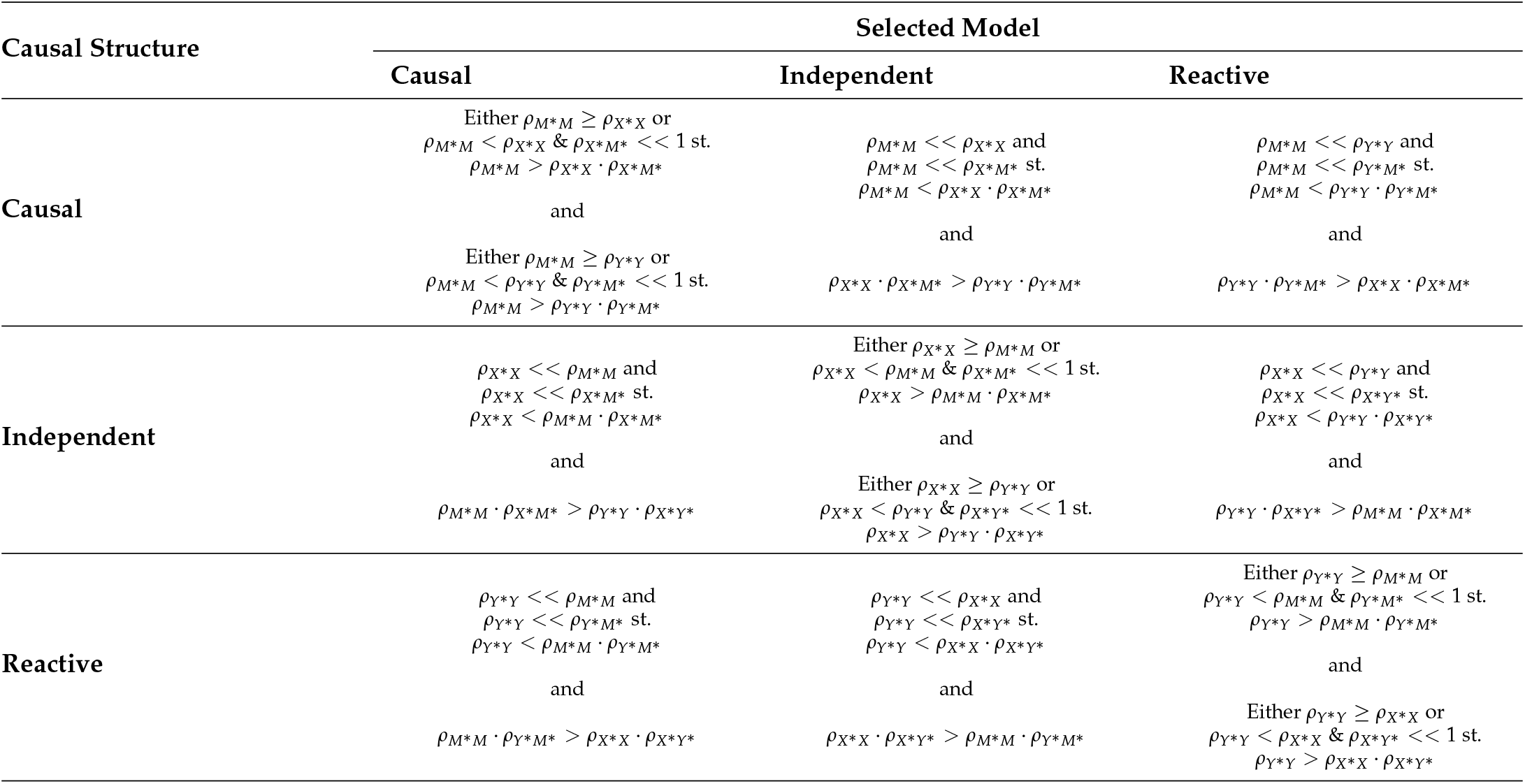
Measurement Error Model Configurations. Configurations of the measurement error model that will lead to consistent inferences (diagonal boxes) versus inconsistent inferences (off-diagonal boxes) when classified with three-choice model options.

